# Using Bayesian priors to overcome non-identifiablility issues in Hidden Markov models

**DOI:** 10.1101/2024.04.20.590387

**Authors:** Jan L. Münch, Ralf Schmauder, Fabian Paul, Michael Habeck

**Affiliations:** Institute of Physiology II, Jena University Hospital, Friedrich Schiller University, Jena 07743, Germany; Department of Biochemistry and Molecular Biology, University of Chicago, Chicago, United States; Microscopic Image Analysis Group, Jena University Hospital, Friedrich Schiller University, Jena 07743, Germany

## Abstract

Hidden Markov models (HMMs) for biomolecules suffer from various forms of parameter non-identifiability. This poses severe challenges to both maximum likelihood and Bayesian inference. However, Bayesian inference offers effective means of overcoming these pathologies. We study the role of prior distributions in the face of practical parameter non-identifiability in Bayesian inference applied to prototypical patch clamp data of ligand-gated ion channels. We advocate the use of minimally informative priors, as they increase the accuracy and decrease the uncertainty of the inference. For complex HMMs, stronger prior assumptions are needed to render the posterior *sufficiently proper*. This can be achieved by confining the parameter space to physically motivated limits. Another beneficial assumption is finite cooperativity of ligand-binding and unbinding events, which introduces a bias towards non-cooperativity but still allows for a non-vanishing degree of cooperativity that is inferred from the data. Despite its vagueness, our prior renders the posterior *sufficiently proper* for all datasets that we considered without imposing the assumption of non-cooperativity. Combining all prior factors allows for meaningful inferences with a dataset of a thousand times lower quality.

## I. INTRODUCTION

Time series data of various processes can be explained by continuous-time Markov Models (MM) [1]. In a biophysical context, the data could probe, e.g., the function of different proteins [2–9] or RNA folding kinetics [10, 11]. Assuming a well-mixed environment, these systems can be modeled by discrete states that interconvert via stochastic transitions, thereby defining a chemical reaction network (CRN). For example, states of a ligand-gated ion channel can be classified by conductance, dwell time, or the number of bound ligands [11–14].

Biophysical experiments typically only observe partial and noisy data such that states with similar signal properties, e.g., conductance, are aggregated into signal classes. Therefore, hidden Markov models (HMMs) must be used to describe the data [15, 16]. A common use case is the analysis of single-molecule ion channel data [17–23] but HMMs are also applied to other experiments [7–11, 24–29].

Often, the mean signal observed in single-molecule and ensemble data is a linear projection of the full Markovian dynamics onto a lower dimensional observable. This can cause the data to be insensitive to the rates of specific subprocesses within the CRN, which complicates their biophysical interpretation. We argue that the dimensionality reduction and aggregation of states, in general, induces a varying degree of practical parameter non-identifiability even for simple CRNs. Experimental noise and limited signal bandwidth only increase the severity of non-identifiability issues. Even worse, the HMM might become structurally non-identifiable [30–34].

Structural non-identifiability refers to models whose parameters cannot be inferred uniquely, even with an infinite amount of data [34]. For example, it might only be possible to infer algebraic combinations of parameters but not the parameters themselves. Instead practical non-identifiability is encountered when there is still a unique optimal parameter set, but it is impossible to collect enough data to reach a sufficiently low parameter uncertainty [33, 35, 36].

Whether a model is structurally or practically non-identifiable depends on the likelihood function, which can be derived from the chemical master equation (CME) in the case of MMs [37]. Here, we study a Bayesian filter [38] based on the Fokker-Planck approximation (FPa) of the CME [39], which preserves the crucial Markov property in a macroscopic signal [38]. The Bayesian filter extends the ideas of Moffat [40] to define a more general and realistic likelihood for ensemble patch clamp (PC) data. A complete HMM inference consists of both parameter estimation and model selection. In some cases, model selection for HMMs can be automated by inferring an infinite HMM [41–43]. However, to the best of our knowledge, the infinite HMM [41–43] only applies to single-molecule data analyzed by discrete-time HMMs but cannot be extended to ensemble data or continuous-time HMMs. Therefore, we assume a fixed CRN topology during parameter inference. Notably, the hidden variable of an HMM does not have to be discrete but can also be continuous [44, 45]. For example, Kalman filters [46, 47] can be used to approximate discrete HMMs [38, 40, 48–51] and define a valid HMM [44, 45].

Parameter estimation via maximum likelihood (ML) [52, 53], profile likelihood [33, 35], maximum a posteriori (MAP), and Bayesian inference [54] suffer in different ways from practical non-identifiability. Limitations in the amount and quality of data (relative to the complexity of the investigated CRN) severely impair or even prohibit ML and MAP inferences [55]. The profile likelihood technique has better uncertainty quantification than ML [33, 35], but still assumes an asymptotic amount of data. A full Bayesian inference [20, 21, 23, 41, 56–59] does not refer to an asymptotic limit in the amount of data and thus has a unique way to deal with parameter non-identifiability. In Bayesian statistics, unknown quantities are treated similarly to random variables [60, 61] in that probabilities express their uncertainty. The prior distribution encodes their uncertainty before analyzing the data. The result of a Bayesian analysis, the posterior distribution, represents the uncertainty of the unknowns in the light of the combined information encoded in the prior and the likelihood. Nevertheless, structural and practical non-identifiability also poses a challenge to Bayesian inference because these pathologies can result in improper posteriors [35, 62].

Our work focuses on the benefits and limitations of minimally informative and vaguely informative priors motivated by physical considerations in the presence of practical non-identifiability. We show that practical non-identifiability can be severely harmful when using uniform priors on the rate matrix of HMMs. In contrast, we suggest using a minimally informative prior [63–66] inspired by Jeffreys [63] and Jaynes [67]. Minimally informative priors are designed to make posteriors as sensitive to the data as possible. We observe that the minimally informative prior increases the accuracy and surprisingly decreases the uncertainty of parameter inferences. Notably, the minimally informative prior will significantly impact the posterior for any plausible amount of data. We explain the origin of these observations by the presence of practical non-identifiability.

Further, we demonstrate that the uniform and minimally informative priors lead to improper posteriors that cannot be normalized due to the practical non-identifiability (in an infinite parameter space [35]). Thus, the minimally informative prior only alleviates the challenges arising from practical non-identifiability of HMMs by making the posterior *sufficiently proper* but does not fully resolve them. A definition of what a *sufficiently proper* posterior means will be given below. We show that rendering the posterior sufficiently proper is the best that can be achieved in HMM inference when using minimally rather than vaguely informative priors. Notably, the same holds for ML inference. Furthermore, the minimally informative prior drastically improves the convergence [68] of the Hamiltonian Monte Carlo sampler [69–73]. Our results show that the limitations of the posterior observed in [35] are due to the use of a uniform prior.

Moving on to more complex HMMs, we show that eliminating the bias from a uniform prior does not solve non-identifiability issues. More information is needed, even if the HMM is structurally identifiable [62], for *sufficiently* proper posteriors. We present two techniques to achieve this. The first option is to enforce theoretically derived upper bounds, such as diffusion limits for binding rates. These restrict the regions in parameter space that contribute to a diverging normalization integral. This often renders the posterior *sufficiently proper*. However, hard theoretical limits are rarely available and might not apply to all parameters that suffer from non-identifiability. As an alternative or as an additional restraint, we suggest coupling each pair of binding rates and unbinding rates softly. These coupling terms bias the CRN towards non-cooperativity without enforcing it. Introducing these vaguely informative priors is more flexible than the common approach of setting parameters equal, which assumes strict non-cooperativity. The vaguely informative prior only defines the scale of plausible positive or negative logarithmic ratios, i.e., cooperativity in homomeric proteins. Hence, our approach infers how likely different degrees of cooperativity are compared to a non-cooperativity bias, which functions as Occam’s razor. The most precise and accurate inferences are obtained if the minimally informative prior is combined with both additional prior assumptions. This combination of prior information allows for meaningful inferences with a thousand times poorer data quality for the most complex HMM that we studied. Notably, this reduction in the data quality that is necessary for meaningful HMM inference is crucial for CRNs of this complexity in the analysis of real-world PC data sets.

## II. PARAMETER NON-IDENTIFIABILITY IN SIMPLE REACTION NETWORKS

Given time series data *𝒴*_*T*_ = *{y*_1_, …, *y*_*T*_*}* of length *T*, and a probabilistic model in the form of a likelihood *p*(*𝒴*_*T*_ |***θ***) := Pr(*𝒴*_*T*_ | ***θ***), the ML approach [74] infers the unknown parameters ***θ***_true_ by maximizing *p*(*𝒴*_*T*_ |***θ***) over the parameter space **Θ**. For models with structurally identifiable parameters, ***θ***_ML_ converges in distribution to ***θ***_true_. The quantification of the uncertainty of the ML estimate ***θ***_ML_ for models that satisfy certain regularity conditions is discussed in Sec. II B. Unfortunately, HMMs do not satisfy these regularity conditions [75]. They are singular instead of regular statistical models. We indicate the possible consequences of singular models by the rate equation (RE) solutions of two toy kinetic models.

### A. Structural parameter non-identifiability

In general, structurally non-identifiable models are characterized by submanifolds in **Θ** in which the likelihood is constant, even with infinitely many data [30, 62]. For the sake of argument, we only look at the RE solution from which an approximate likelihood can be derived. An example of a structurally non-identifiable model is a linear birth-death process characterized only by the mean number of bacteria 𝔼 [*n*_bakt_(*t*)] in a well-stirred petri dish:

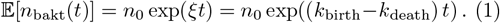

Parameter pairs ***θ*** = (*k*_birth_, *k*_death_)^*⊤*^ with constant difference *ξ*(*k*_birth_, *k*_death_) = *k*_birth_ *− k*_death_ result in the same 𝔼 [*n*_bakt_(*t*)]. This model is structurally non-identifiable, because one cannot disentangle *k*_birth_ and *k*_death_ based on 𝔼 [*n*_bakt_(*t*)] alone. A likelihood derived only from 𝔼 [*n*_bakt_(*t*)] would show the symmetry *p*(*𝒴*_*T*_ |***θ***) = *p*(*𝒴*_*T*_ |*ξ*). This implies that the likelihood is flat, *p*(*𝒴*_*T*_ |***θ***) = const, along straight lines in **Θ** with intercept *ξ*. Hence, the ML estimator cannot converge to a normal distribution centered at ***θ***_true_. However, there is independent information in the higher-order statistical moments that renders the linear birth-death model structurally identifiable. By incorporating the information contained in var[*n*_bakt_(*t*)], which can be derived from the CME [76], one obtains a structurally identifiable model. Thus, structural non-identifiability can be caused by ignoring higher statistical moments of the data-generating process. Similarily, ignoring the Markov property of equilibrium fluctuations leads to structural non-identifiability in HMM inference as shown in [40].

### B. Practical parameter non-identifiabilty

Structural identifiability is necessary, but not sufficient for successful ML inferences. For a finite amount of data, the HMM must also be practically identifiable, or, as we prefer to argue, it must be *sufficiently* practically identifiable. Let us clarify what we mean by this. The literature offers different definitions of practical identifiability vs. practical non-identifiability. Here, we follow the definitions of [33]. Likelihoods suffering from practical non-identifiability do not decay to zero (Fig. 1, blue curve for *θ > θ*_max_), but stretch out infinitely in regions of **Θ** (in one or multiple dimensions) for any finite amount of data [33]. This happens already in simple, partially observed CRNs.

**FIG. 1.**
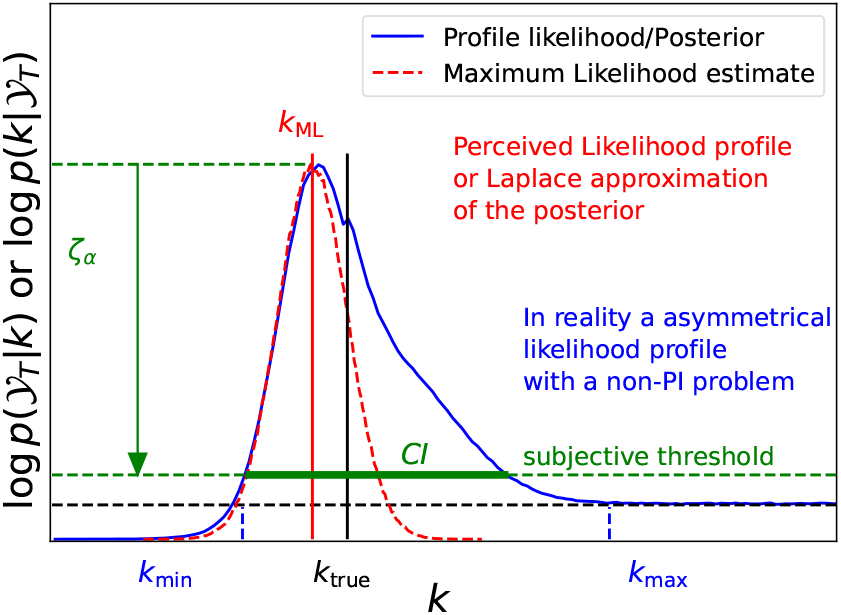
The severity of practical parameter non-identifiability depends on the relative height of the peak compared to the non-vanishing tails. We sketch a one dimensional inference problem, e.g., a unknown chemical rate *k*. The red dashed line shows the ML inference or Laplace approximation of the posterior based on a uniform prior. Note that the ML inference (using the curvature) typically underscores the uncertainty (quicker decay of the red dashed line than the blue solid line), particularly for *k > kML*. Note that a prediction of the uncertainty based on the curvature at *k*_ML_ cannot detect these shortcomings, because any function can be approximated with a second-order Taylor expansion around extreme values.

Let us assume that we can record the occupation number *n*_*O*_(*t*) of state **O** at any frequency and without any measurement noise such that, for the sake of argument, the additional complications arising from noisy data are avoided. As an example for a practically non-identifiable likelihood consider a CRN with a rate *k*_BA_ · *L* that depends on ligand concentration *L* (or any other stimulus-dependent rate):

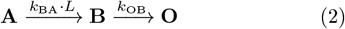

for a finite number of channels. Assuming that the initial condition is **n**(*t*_0_) = (*n*_*A*_, 0, 0) and that the experimental readout is *y*(*t*) *∝* 𝔼 [*n*_*O*_(*t*)], the part of the general solution of the RE that is experimentally accessible is

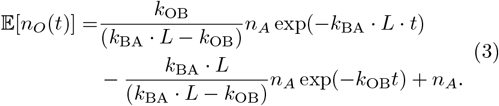

Note that the solution of the CME for the initial condition **n**(*t* = 0) = (*n*_*A*_, 0, 0) is a multinomial distribution. To understand this, consider the case that only one molecule is in state **A** at *t* = 0. For all *t >* 0, the molecule has probability *p*_*O*_(*t*) to be in **O**, *p*_*B*_(*t*) to be in **B** and *p*_*A*_(*t*) to remain in **A**. If, instead, one has *n*_*A*_ independent and identical molecules in state **A** at *t* = 0, then each of them individually has the same probability *p*_*O*_(*t*), *p*_*B*_(*t*) and *p*_*A*_(*t*) at *t >* 0. Hence, the distribution over all states is a multinomial distribution [77] that evolves over time. If only state **O** is observed, one can reduce the problem to

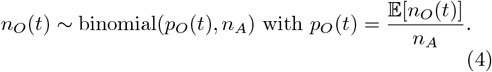

If none of the rates could be changed externally by varying *L* such that *k*_BA_(*L*) = const, then we would also face structural non-identifiability. For a ligand-dependent rate *k*_*BA*_ · *L*, we can run the experiments at different ligand concentrations *L* and thereby overcome structural non-identifiability. However, if we measure at two concentrations *L*_1_, *L*_2_ such that *k*_*BA*_ · *L*_1_ and *k*_*BA*_ · *L*_2_ and *k*_*OB*_ are all different, but similar in magnitude, then we can still face a practical non-identifiability problem. Practical parameter non-identifiability originates from the following phenomenon: Any combination of values for *k*_BA_ · *L*_*j*_ and *k*_OB_ that satisfies

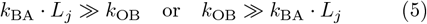

will push the amplitude of one of the two exponential decays (Eq. 3) to zero. Even if experiments were run at many different ligand concentrations *L*_*j*_, we could still find combinations of *k*_BA_ and *k*_OB_ that satisfy one of the conditions in Eq. 5 for all *L*_*j*_. In regions of **Θ** where one of the conditions (Eq. 5) holds for all *L*_*j*_, minor changes in *k*_BA_ or *k*_OB_ will hardly affect 𝔼 [*n*_*O*_(*t*)]. Note that the correct solution of the CME is a multinomial distribution given the initial conditions. Adding information about the entire distribution of *n*_*O*_(*t*) such as variance or skewness, etc. will not resolve the practical non-identifiability problem caused by the vanishing amplitude of one of the exponential decays. However, the multinomial model would improve the accuracy and quality of the uncertainty quantification.

This example is reminiscent of the common scenario for coupled CRNs in which only *k*_*j*_ can be inferred if *k*_*i*,true_ ≫ *k*_*j*,true_. However, here we do not consider a scenario in which the signal of the data-generating process is rate-limited. Instead, we assume that at least three different rates are at play, *k*_BA_(*L*_1_), *k*_BA_(*L*_2_), and *k*_OB_ that have similar magnitude but non-identical values. Therefore, rate-limiting contributions do not exist at least for one combination of *k*_BA_(*L*_*i*_) and *k*_OB_ in the data-generating process. Nevertheless there are regions in parameter space **Θ**, far away from the true parameter values ***θ***_true_, in which one or the other rate is rate-limiting for the predictions of the model. The structure of the CRN together with the fact that the signal is generated by a linear projection have the potential to be rate-limited somewhere in **Θ**, independent of the true parameters of the data-generating process ***θ***_true_. Thus, for models such as *n*_*O*_(*t*) ∼ binomial(*p*_*O*_(*t*), *n*_*A*_(0)), the likelihood will approach a non-vanishing constant due to rate-limiting effects in regions where *k*_*i*_ ≫ *k*_*j*_ and hence become practically non-identifiable. See App. A for a discussion on the effects if states **A, B** are observed in isolation or simultaneously.

Structural identifiability is a binary property (a model is either structural identifiable or not), whereas practical identifiability is gradual (continuous) [78]. The likelihood function (Fig. 1 blue curve) which is proportional to the posterior *p*(*k*|*𝒴* _*T*_) (for a uniform prior) indicates the continuous nature of practical non-identifiability. The maximum value of *p*(*𝒴*_*T*_ | *k*) relative to the constant value in the tails specifies the degree of parameter identifiability. However, to classify models into practically identifiable or practically non-identifiable, one uses the confidence interval CI based on confidence level *ζ*_*α*_:

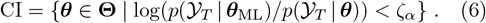

One defines the model as practical identifiabile if the confidence interval CI is a compact set. This holds if the subjectively defined threshold *ζ*_*α*_ (Fig. 1) [76] is larger than the asymptotic value of the likelihood *p*(*𝒴*_*T*_ |***θ***_flat_) (Fig. 1, black dashed line). For multi-parametric models, there will be multiple asymptotic values for the different directions in **Θ**. The interval [*k*_min_, *k*_max_] where the dashed black lines cross the likelihood profile (blue lines) (Fig. 1) is the largest asymmetric confidence interval that can be deduced by using the profile likelihood technique [33]. The data contain no information to distinguish values of *k* larger than *k*_max_. However, the data are informative for values *k < k*_max_ and even *k < k*_min_. Often, the profile of *p*(*𝒴*_*T*_| ***θ***) might approach a non-vanishing constant value only asymptotically. Subjectively defining a threshold relative to the maximum value of the likelihood [33] is equivalent to choosing a significance level []. Note that ML does not infer the shape of *p*(*𝒴*_*T*_ |***θ***) globally (Fig. 1 red dashed curve). ML estimates the shape based on the curvature of the likelihood at ***θ***_ML_ using the Fréchet-Darmois-Cramér-Rao bound theorem. Therefore, standard ML does not detect practical non-identifiability (Fig. 1 red dashed curve). The degree to which practical non-identifiability affects the parameter inference depends on, as discussed, the intrinsic properties of the MM (i.e. the number of states and their connectivity) and on the specifics of the experimental data such as the rank of the linear projection, the signal-to-noise ratio and the signal bandwidth.

Due to the challenges indicated by the two toy examples, HMM inference based on ML has several drawbacks compared to sampling from the posterior *p*(***θ***| *𝒴*_*T*_). First, one must take extra precautions against pathologies of *p*(*𝒴*_*T*_ |***θ***) resulting in structual or practical non-identifiability, because ML does not reveal them [33]. Second, even if the model is structurally identifiable and sufficiently practically identifiable, the quality and quantity of the data are often insufficient to meet the implicit assumption that *p*(***θ***_ML_) can be approximated by a normal distribution, which is a requirement to justify the use of ML. For a comment on strategies to detect parameter non-identifiability see App. 1. Fortunately, Bayesian statistics can deal with these pathologies of the likelihood. Nevertheless structural and practical non-identifiability pose a challenge. They create regions in **Θ** where the prior dominates entirely the likelihood.

## III. BAYESIAN INFERENCE IN A NUTSHELL

The Bayesian posterior

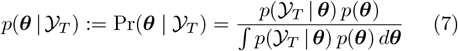

is a probability distribution on **Θ** and combines the information encoded in the prior *p*(***θ***) := Pr(***θ***) and the likelihood to quantify the uncertainty of ***θ***. The posterior is called “proper”, if it can be normalized, which means the denominator in (Eq. 7) satisfies

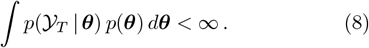

We introduce the terminology *sufficiently proper* to clarify the notion of practical parameter non-identifiability. Practical non-identifiability in combination with minimally informative priors, which are often improper, may result in posteriors that are improper in a strict sense. The blue curve in Fig. 1 illustrates this for a one-dimensional case and an improper uniform prior. The essential information making the posterior proper (Fig. 1) are the inconspicuous cutoff values of the uniform prior.

The higher the posterior the less sensitive is the inference to actual values of the cutoff. Thus, we define the ratio of the height of the posterior (based on a uniform prior) at the MAP estimate and its non-vanishing asymptotic value as

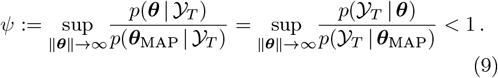

One could subjectively define the posterior to be *sufficiently proper*, if *ψ ≤* 10^*−*3^, say, meaning that the peak of the posterior/likelihood towers at least by three orders of magnitude over the flat parts of the likelihood (in all unbounded directions in **Θ**). For *ψ* = 0, the posterior is sufficiently proper and might even be strictly proper. For non-uniform priors and one-dimensional inferences, flat parts of the likelihood reveal themselves by a posterior that is proportional to the prior in that area.

In multi-parameter inference problems with non-uniform priors the situation is in general more complicated. The likelihood could approach a non-vanishing constant value on some unbounded subset **Θ**_flat_ *⊂* **Θ** of any shape. However, we will encounter the simpler case, that marginal posteriors are proportional to the prior in certain areas. This can be explained by assuming that the likelihood is flat for an unbounded set in the direction of parameter *θ*_*i*_. Let us denote the parameters without *θ*_*i*_ by ***θ****\*_*i*_ and also assume that the prior factorizes: *p*(***θ***) = *p*(***θ*** *\*_*i*_) *p*(*θ*_*i*_). Then

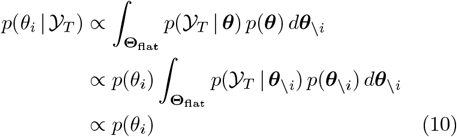

holds for the marginal posterior locally. So changes in the posterior are proportional to changes in the prior along *θ*_*i*_ in that region of the parameter space. If **Θ**_flat_ is more complicated (any differentiable curve or hyperplane), then one need to check if changes in the posterior are proportional to changes in the prior, when moving within **Θ**_flat_. From a practical perspective, we call a posterior to be sufficiently proper, if *ψ* (for a posterior based on a uniform prior) is small enough such that sampling from the posterior of interest (based on the minimally informative prior) is insensitive to moderate changes in the limits of the sampling box. Only if these limits are increased by orders of magnitude, then the posterior is going to be affected in its lower-order statistical moments. We will see that minimally informative priors, while sensitizing the posterior to the data, desensitize the resulting posterior to the exact limits of the sampling box and thereby render posterior sampling more robust. That way, it will become possible to analyze a dataset with, i.e., *ψ* = 0.05, if we use a minimally informative prior. We will also demonstrate that parameter inference can be improved further if we include additional information via the prior.

We will also use a simpler definition of *sufficiently proper* based on posterior samples, namely if the posterior mode carries most of the probability mass such that samples from the tail region hardly reach the limits of the sampling box. This indicates that probability mass in the tail regions is negligible relative to the probability mass under the posterior mode. Note that the density is only well-defined because we refer to a finite volume in the parameter space, otherwise the posterior is improper.

We refer to an inference as fully Bayesian if Eq. 7 is calculated or sampled

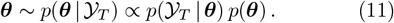

We use Hamiltonian Monte Carlo (HMC) [69, 70, 73] as provided by the Stan software [71, 72] to generate samples from *p*(***θ***| *𝒴*_*T*_). In addition to the covariance matrix of the parameters, *p*(***θ***| *𝒴* _*T*_) allows the calculation of the credibility volume in order to assess parameter uncertainty. The smallest volume *V*_*P*_ that encloses a probability mass *P ∈* [0, 1]

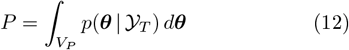

is called the Highest Density Credibility Volume (HDCV). Assuming that the model sufficiently captures the data generating process, the true parameter values ***θ***_true_ will lie in the HDCV with a probability *P* as soon as the likelihood dominates the prior.

Bayesian inference is conditional on the assumed prior and likelihood [79]. Altering *p*(*𝒴*_*T*_ |***θ***) or *p*(***θ***) changes *p*(***θ***|*𝒴* _*T*_). The prior becomes irrelevant only in the infinite data limit (and only for regular models), meaning that ML and Bayesian inference become equivalent [54]. In case of practically non-identifiable models, Bayesian inference has at least two advantages over ML. First, by scrutinizing *p*(***θ*** | *𝒴*_*T*_) in detail, issues with structural or practical non-identifiability are revealed. Second, the introduction of priors can alleviate non-identifiability problems.

If little is known *a priori* about reasonable parameter values and the data constrain some parameters only vaguely, the use of a minimally informative prior is essential. It attempts to make *p*(***θ*** |*𝒴*_*T*_) as sensitive to the data as possible. Typically, a minimally informative prior maximizes the variance of the posterior *p*(***θ*** |*𝒴*_*T*_). In contrast, we show that the minimally informative priors introduced below help confine *p*(***θ*** |*𝒴*_*T*_) within reasonable boundaries. Thus they reduce the variance of the posterior, if compared to uniform priors. However, minimally informative priors themselves are often improper, which might also render *p*(***θ*** | *𝒴*_*T*_) improper if *p*(*𝒴*_*T*_| ***θ***) is practically or even structurally non-identifiable. Thus, the posterior *p*(***θ*** |*𝒴*_*T*_) will be dominated by the prior *p*(***θ***) in regions of **Θ** where data fail to inform us about the parameters. Fortunately, Bayesian statistics provides us with tools to render *p*(***θ*** |*𝒴*_*T*_) sufficiently proper such as theoretically derived upper limits on parameters or vaguely informative assumptions about cooperativity incorporated in *p*(***θ***). The benefits and limits of combinations of minimally and vaguely informative priors in the presence of practical non-identifiability will be discussed for two CRNs (Fig. 2).

**FIG. 2.**
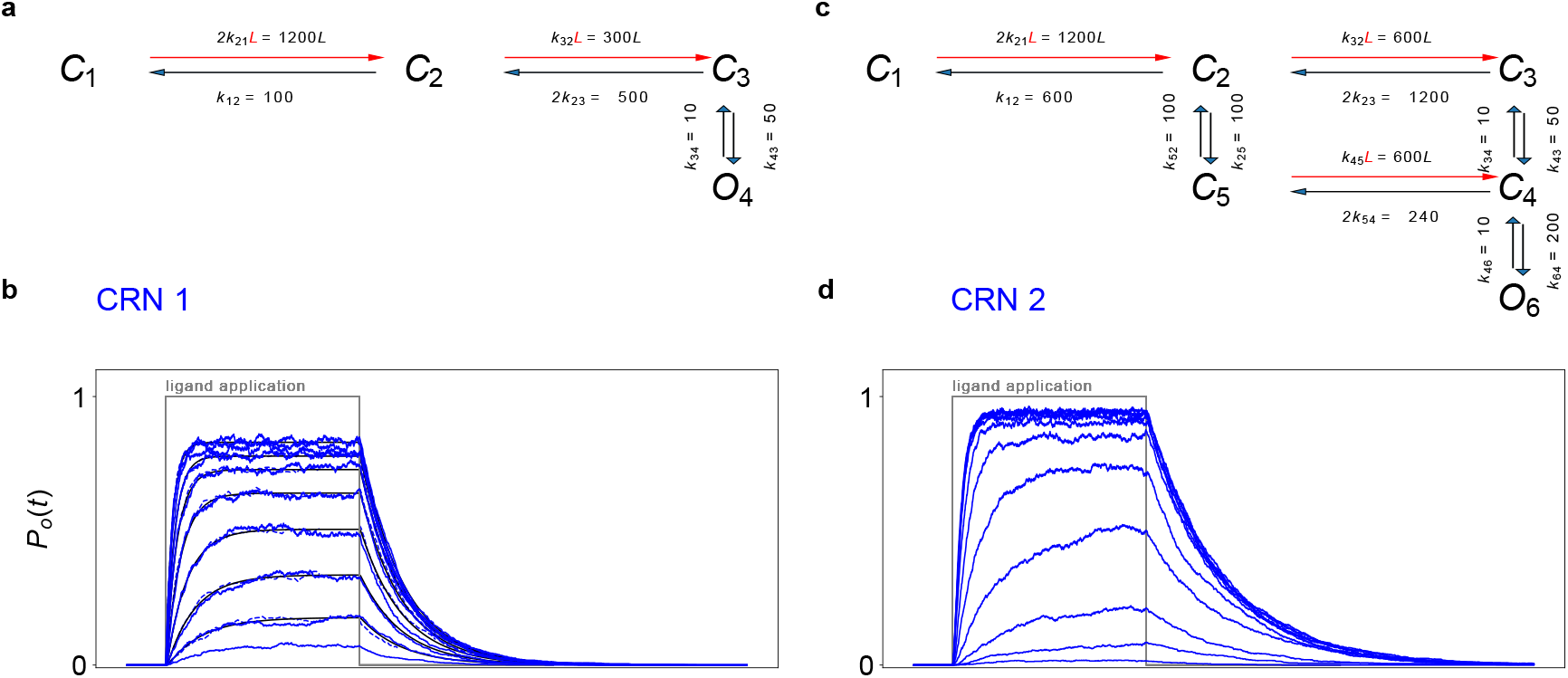
Chemical reaction networks of ligand-gated ion channels and their simulated patch-clamp data. **a, c** The MMs (one MM per column) consist of three or five closed states “*C*_*j*_ “ and one open state “*O*_*j*_ “. Binding steps (red arrows) have concentration-dependent rates. The CRNs are specified by the absolute rate constants *k*_*ij*_, and *L*, the ligand concentration. Rates are given per subunit. Stoichiometry factors account for the number of subunits able to undergo the respective transitions. The units of the rates are in a. u. To calculate their SI units s^*−*1^ and *µ*M^*−*1^s^*−*1^, one needs to multiply their value by 6*/*7 (Sec. V C 4). The open states *O*_*j*_ conduct a mean single-channel current *i* = 1, and the closed states *C*_*j*_ conduct a current of *i* = 0. The synthetic data were simulated with the Gillespie algorithm at a sampling rate of 10 ka.u. The Bayesian filter analysis frequency *f*_ana_ is 2 to 5 ka.u. Since the units are in a. u., the ratios of rates or inverse dwell times determine the CRN. Further their ratio to the sampling frequencies determines how detailed the kinetics are recorded. Similarly, the relative magnitude of *i* compared with *σ*_op_ and *σ*_ex_ should be used to relate simulations to experimental conditions. **b, d** Open probability time traces calculated from normalized currents 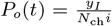 of simulated relaxation experiments of ligand concentration jumps with *N*_ch_ = 10^3^ channels. For demonstration purposes, no experimental noise is added in this figure such that all fluctuations originate from Markov state transitions. However, when inferring the posteriors, additional experimental noise is added. The black lines are the theoretical open probabilities *P*_o_(*t*) of the model. Typically, we used the set {0.0625, 0.125, 0.25, 0.5, 1, 2, 4, 8, 16, 64} *µ*M of 10 ligand concentrations.

## IV. PARAMETRIZATION OF THE RATE MATRIX

In the following, we will analyze patch-clamp data that are simulated with MMs involving only mono-molecular or pseudo-monomolecular chemical reactions such as conformation dynamics of a protein or binding/unbinding transitions at excess ligand. If the CRN describes a single molecule, it can only be in one of *M* Markov states at time *t*:

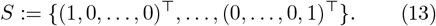

where we use a one-hot encoding of states. If **s**(*t*) = **e**_*i*_ where 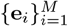 denotes the standard basis, then the channel is in state *i* at time *t*. State-to-state transitions are governed by a transition matrix **T** in discrete time,

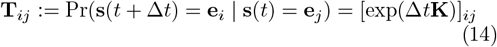

with time increment Δ*t*, or with a rate matrix **K** in continuous time:

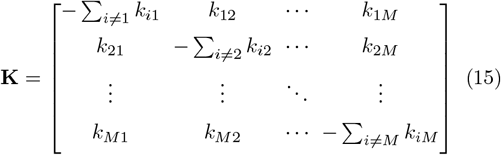

where *k*_*ij*_ *≥* 0 for *i≠ j*. By definition, each column of **K** sums to zero reflecting the assumption that the CRN is closed. The dwell times to remain in the *i*-th state are exponentially distributed with mean 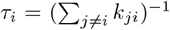. To facilitate the definition of a minimally informative prior in the next section, we use an alternative parameterization of **K** that does not involve the chemical rates *k*_*ij*_:

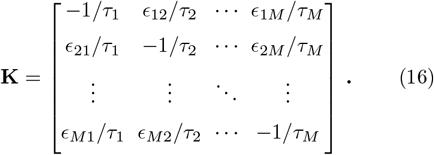

The parameters *ϵ*_*ij*_ denote the probability of transitioning from state *j* to state *i* after the random dwell time has passed. Thus, each chemical rate 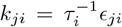 is the product of a probability with the inverse mean dwell time. The parameters *ϵ*_*ji*_ have no units, unless the statistical weight corresponds to a ligand-dependent *k*_*ji*_. Since ∑_*j*_ *ϵ*_*ji*_ = 1, both parameterizations of **K** have the same number of free parameters. For each column, we separated the inverse time-like scale parameters 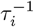 from the probabilities *ϵ*_*ji*_, which are shape parameters. Because transition probabilities are constrained to 0 *≤ ϵ*_*ji*_ *≤* 1, the likelihood remains finite for all *ϵ*_*ji*_ and *p*(**K** |*𝒴*_*T*_) will be proper in these parameters (as long as Haldane-like priors beta(0, 0) are excluded for *ϵ*_*ij*_). Then, only the dwell time parameters *τ*_*i*_ can render the HMM practically non-identifiable.

Figure 2 shows the time traces of plausible CRNs of two ligand-gated ion channels that have two binding pockets. These can be simulated with QuB [80] or an inhouse algorithm https://cloudhsm.it-dlz.de/s/QB2pQQ7ycMXEitE. The assumptions made to define a likelihood for these data are detailed in [38]. In App. B, we discuss the global sensitivity of the solution of the RE for CRN1 (Fig. 2 **a**) and demonstrate practical non-identifiability of the likelihood (App. B 3) even for over-optimistic data and strong prior knowledge (only a single rate constant is unknown).

## V. DEFINING AND BENCHMARKING THE MINIMALLY INFORMATIVE PRIOR

The following section benchmarks the performance of the Bayesian filter for different combinations of minimally informative priors and physically motivated vaguely informative priors, for cases where the information content of the data is low relative to the number of parameters of the CRN. We first compare a uniform prior on the rate matrix *p*(**K**) with a minimally informative prior defined below (Eq. 20), which promises to be less biased (less misinformed). The reason is that the practical non-identifiability of the likelihood (App. B 1) is aggravated when using a uniform *p*(**K**). In an unfortunate combination, a uniform prior *p*(**K**) places the probability mass where the likelihood becomes less and less pronounced and reaches a constant finite value (App. B 1). With a minimally informative prior, however, one can drastically reduce the severity of this problem. But one should be aware of the limitations of both priors. See App. C 1 for a brief biophysical example for the problem of different parametrizations of statistical models and prior distributions, which gave rise to the following Eq. 17.

### A. Definition of minimally informative (MI) priors by approximating Jeffreys’s rule

We will use a revised version of Jeffreys’s rule [64] to define the minimally informative prior, which treats location, scale and shape parameters independently:

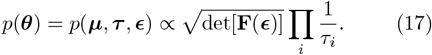

The location parameters, ***µ***, such as the mean value of the normal distribution, are assigned uniform priors. Each scaling parameter, ***τ***, has a log-uniform prior. Only the shape parameters ***ϵ*** are treated conjointly by evaluating the Fisher matrix,

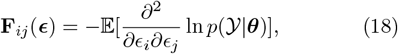

according to Jeffreys’s rule [64]. The separated treatment of location, scale and shape parameters can be applied to the used parametrization (Eq. 16) of **K**. For a brief introduction of Eq. 17, see App. C 2. In addition, we simplify Eq.17 by assuming that 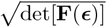 can be applied to each column of **K** (Eq. 16) independently. In that way, we obtain closed-form solutions of 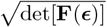 derived from simpler statistical models for the remaining ***ϵ*** of each column of **K**.

### B. Minimally informative prior for the rate matrix inspired by Jeffreys’s rule

We use a simplification of Eq. 17 that is common practice [11] for complex multi-parameter MMs. The priors used to infer MMs from MD simulations are constructed by applying Jeffreys’s rule to simpler statistical models [11] instead of applying it to the entire model. Bayesian estimation of **T** (Eq. 14) from MD simulations [81] often uses one Dirichlet prior per column **T**_*i*,:_ of the transition matrix:

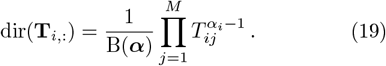

However, the Jeffreys prior for **T** is not a product of Dirichlet distributions [82, 83]. Also, for HMMs of single-molecule force spectroscopy data [10], products of Dirichlet priors are used for **T**.

Here, we do not sample **T** but **K** (Eq. 14) because, in contrast to sampling **T**, it is trivial to incorporate information about the scaling of the binding rates at different ligand concentrations in a direct parameterization of **K**. The same applies to additional prior information on maximal binding rates. Note that with one exception, we use the parameterization of Eq. 16 to define *p*(**K**). The exception will be discussed later when we add the information about theoretical upper diffusion limits on binding rates *k*_21_ and *k*_32_ (CRN1 and CRN2) and *k*_45_ (CRN2). One can mix in the parameterization any *k*_*ji*_, *τ*_*i*_ and *ϵ*_*ji*_ equivalently as long as the pior remains equivalent, i.e., that 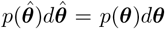 holds, with 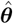 indicating a different parameterization. As the mean dwell times *τ*_*i*_ are scaling parameters, we use a log-uniform prior following the arguments above. We use the Dirichlet prior for the probabilities *ϵ*_*ji*_, which is the default prior for probability vectors. These probabilities should not be confused with the transition probabilities stored in **T**. Applying Eq. C3 to a multinomial likelihood, results in a Dirichlet prior with *α*_*j*_ = 0.5 for all *j ∈* [1, · · ·, *W*_*i*_] parameters. The value of *W*_*i*_ is the number of Markov transitions leaving the *i*-th state. For a physically meaningful CRN, **K** tends to be sparse, e.g., ligand binding and channel gating does not occur at the exact same instant of time. Thus for the *i*-th Markov state, the number of allowed transitions will usually satisfy *W*_*i*_ *< M* where *M* is the total number of states. The set Ω_*i*_ *⊆{*1, …, *M}*/ *{i}* contains the indices of all states that can be reached by one Markov transition, leaving the *i*-th state. Then, using Eq. 17 and evaluating the 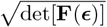 factor for each column of **K** individually, we obtain

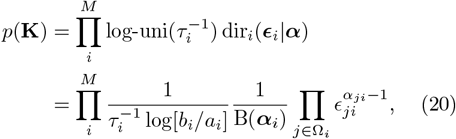

where ***α*** is the vector of concentration parameters and B(***α***) is the beta function at ***α***. The set of *a*_*i*_ and *b*_*i*_ for the log uniform distributions are the upper and lower limits of *τ*_*i*_. The topology of the CRN is controlled by the Dirichlet distribution. See App. D for an illustration of *p*(**K**) for a state with two or three leaving transitions.

### C. Advantages and limitations of the minimally informative prior in the presence of practical non-identifiability

Using the Bayesian filter, we study the impact of three different priors on the performance of *p*(**K**| *𝒴*_*T*_), focusing on weakly informative patch-clamp data. The data are sampled from a 4-states-1-conducting-state HMM (CRN1 Fig. 2**a**). Our findings are presented in Fig. 3.

**FIG. 3.**
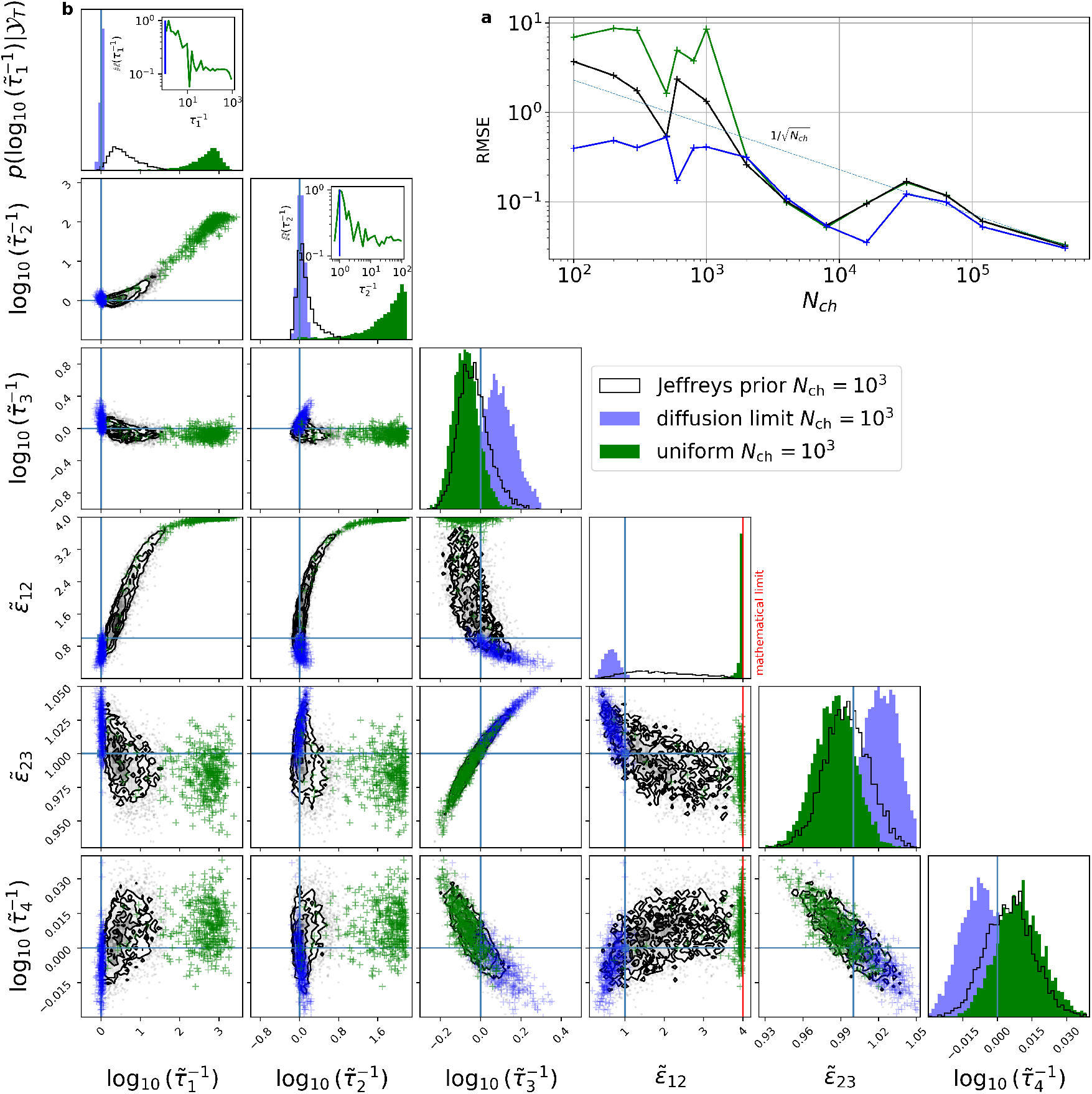
Prior sensitivity analysis showing that the uniform prior aggravates problems caused by practical non-identifiability. The figure contrasts the posteriors resulting from three different priors: uniform prior (green), minimally informative prior (black), and minimally informative prior with physical limits on the binding rates (blue). The rates are transformed to inverse dwell times and transition probabilities. **a**, RMSE of the mean of the marginal distribution vs. *N*_ch_ based on the uniform prior (green), the minimally informative prior(black) and minimally informative prior with imposed imposed diffusion limits (blue). The cyan dashed curve is a fit based on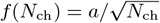. A standard deviation of each data point which follows 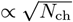 is assumed. All data sets were generated with *σ*^2^ = 1 and 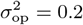. **b**, Posterior distributions of the dwell times *τ*_*i*_ and transition probabilities *ϵ*_*ji*_ for *N*_ch_ = 10^3^. On the diagonal, 1D marginal posteriors are plotted. The off-diagonal plots show the 2D marginal distributions of the posterior. All samples of the parameters were normalized to their true values 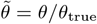. The blue lines indicate the true parameters. The insets on the diagonal show the same histogram only plotted on the logarithmic scale to display the flatness of the posterior if a uniform prior is used. The (red) vertical bar indicates *ϵ*_12_ = 1, which corresponds to 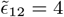 due to the normalization by the true values. The posterior is plotted based on samples with a Gelman-Rubin statistic of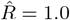.

#### 1. The scaling of the RMSE

We define the Euclidean distance or root-mean-square error (RMSE) between **K**_true_ and the posterior mean of all chemical rates in log-space as

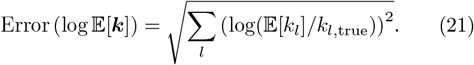

Appendix E discusses why we use the logarithm of the chemical rates. In Fig. 3**a**, the RMSE of the inferred *p*(**K**|*𝒴* _*T*_) is plotted against the number of ion channels per time trace *N*_ch_. We use *N*_ch_ as a proxy for the quality or information content of the data. A regular statistical model is expected to show a 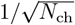-scaling of the RMSE, but the Bayesian filter is singular. This becomes apparent for the uniform prior: Below a critical value *N*_ch,crit_ ≈ 2·10^3^ the RMSE deviates visually from the behavior of the other two priors. Above *N*_ch,crit_ the Bayesian filter behaves like a regular statistical model.

In the biased regime, *N*_ch_ *< N*_ch,crit_, the uniform prior causes the RMSE to deviate from the 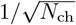-scaling, whereas the log-uniform prior scales as 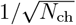 to far smaller *N*_ch_, because it imposes a penalty on larger values of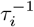. Deviations from 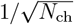-scaling of the RMSE to smaller values also occur if upper bounds are enforced on some *k*_*ij*_. The log transformation eliminates the lower limits (App. E), but the upper bounds derived from the diffusion limit (Sec. V C 4) are still active. In contrast to the equivalence of the minimally informative and uniform prior above *N*_ch,crit_ the parameter limits reduce the RMSE even at *N*_ch_ that are two orders of magnitude larger than *N*_ch,crit_. Note that the RMSE is dominated by the error in 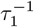whose uncertainty is reduced most strongly by the constraints (Fig. 14). The impact of the other upper limits is discussed below.

#### 2. The likelihood dominates the posterior only in a small region of the parameter space

Next we explore the impact of the prior in more detail for a dataset in the critical regime (*N*_ch_ = 10^3^). The symbol 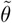 denotes a parameter divided by its true value such that 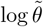should ideally be zero. Figure 3**b** shows the marginal posteriors of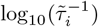. With a uniform prior, the posterior of 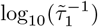 concentrates between 10^2^ and 10^3^ and would deviate even more strongly from zero, if the sampling box was larger. The insets in the diagonal panels provide an overview of the marginal posteriors of 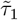 and 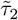over the full sampling box. To indicate the relative probability masses, the posterior ratios 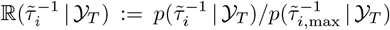 with *i* = 1, 2 are plotted. The flat right tail of the marginal posterior obtained with the uniform prior (Fig. 3**b**, green curve) causes the observed deviations of the RMSE (Fig. 3**a**, green curve and Fig. 13). Furthermore, a flat prior-dominated part of 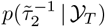 creates an exponentially growing part in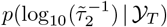. The ratios 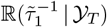 and 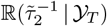 drop from their peak values to their right tails only by *≈* 1*/*10 before they reach a non-vanishing plateau. It takes 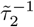 two orders and 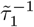 three order of magnitude to do so. Hence, the posteriors 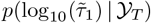 and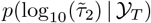, are dominated by the uniform prior. It is plausible that this also holds for the entire *p*(**K**| *𝒴*_*T*_).

Given that the sampling box covers multiple orders of magnitude for 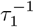 and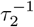, the flat part of the marginal posterior (where the data provide no information about these parameters) contributes a non-negligible probability mass to the posterior. Thus, posterior statistics such as mean, median, variance and derived quantities such as the RMSE become sensitive to the probability mass residing in the flat part. That should raise one’s concern, because the inference will depend on the very limits of the sampling box. A larger sampling box would move probability mass from the peak into the flat area. Hence, when using a uniform prior in the critical regime, the limits of the sampling box for 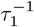 and 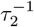 act as highly informative user settings, even though they are often chosen arbitrarily and lack a physical justification. If instead 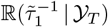 were to drop to 10^*−*4^, say, before the marginal posterior flattens out, changes in the limits would not affect *p*(**K**| *𝒴*_*T*_), unless one increased the sampling box by many orders of magnitude. Because the posterior of *τ*_*i*_ is improper, we cannot define meaningful HDCVs to quantify their uncertainty, if *τ*_*i*_ is defined on the entire real axis. More information is needed, which unfortunately happens often by setting the sampling box limits more or less arbitrarily. Alternatively, vaguely informative priors can be defined by using physical justified information that penalize the tails of *p*(**K** | *𝒴*_*T*_) as demonstrated below.

#### 3. The minimally informative prior (partially) alleviates practical non-identifiability

A comparison of the posteriors obtained with a uniform and the minimally informative prior exemplifies the harm induced by the uniform prior (Fig. 3, black vs. green curve). The minimally informative prior (Eq. 20) penalizes large *τ*_*i*_ and decorrelates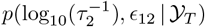. Note that 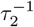 and *ϵ*_12_ belong to the same column in **K** describing transitions leaving state *C*_2_. This accumulates probability mass at the true parameter values in 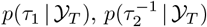and *p*(*ϵ*_12_ | *𝒴*_*T*_). Consistently, the choice of prior has a stronger impact on *τ*_1_, *τ*_2_ and *ϵ*_12_ (which are much less determined by the data) than on 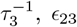 and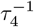. Overall, the minimally informative prior concentrates *p*(**K**| *𝒴*_*T*_) closer to **K**_true_.

Thus, when looking at the RMSE as a function of *N*_ch_, the minimally informative prior reduces the RMSE drastically below *N* = 2 · 10^3^ (Fig. 3**a**) such that the 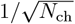 regime expands. The 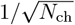 scaling is unfortunate if one tries to increase the inference quality by measuring additional data. However, the 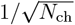 scaling is desirable in the reverse direction in which *N*_ch_ decreases, because it prevents inferences from becoming to quickly meaningless. Despite its heuristic definition, the prolonged 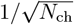-regime indicates that with this minimally informative prior one is less biased than with the uniform prior. Below *N*_ch,crit_ one only sees the prior and hardly any effect of the data when using the uniform prior. The area around the true values has essentially no probability mass. However, the minimally informative prior concentrates the probability mass much closer to **K**_true_. Nevertheless, if the likelihood is flat in 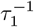 and 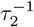 at some distance to the peak, it follows that 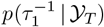 and 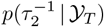are improper. This does not change for the minimally informative prior, because the log-uniform prior cannot be normalized when defined over the entire positive real axis. It diverges at *τ*_*i*_ *→*0 and decays too slowly to zero for *τ*_*i*_ *→ ∞*. We return to this observation in sec. V D.

Cutting **Θ** with a sampling box into regions that are accessible and inaccessible to the sampler is not always a problem. Without knowing about practical non-identifiability problems, we showed in [38] that the HDCVs and thus the estimator have a frequentist interpretation as long as the peak of the posterior is towering highly enough over the flat parts. Thus, one finds **K**_true_ inside a volume with a certain probability mass with a frequency approximately equal to that probability [38].

#### 4. Adding information from physically motivated upper limits for binding rates

So far, we have shown that using a minimally informative prior robustifies the inference below *N*_ch,crit_. To alleviate the problem that the posterior could still be improper or vague in some parameters, we can add more physically motivated information, such as the diffusion limit for the binding rates (Fig. 2, red arrows). Typically, the random collision rate is used as a upper limit for binding rates [84]. Here we use a more realistic estimate for the binding of small ligands [85]. To the best of our knowledge, only Bayesian statistics can rigorously take full advantage of limits on the parameters, because the introduction of parameter limits will impair the validity of the normal approximation of the sampling distribution of ***θ***_ML_. To implement the diffusion limit consistently in a minimally informative way, we need to change the definition of the prior Eq. 20. The rate holds *k*_32_ = *ϵ*_32_*/τ*_2_ such that we can also draw *k*_32_ ∼ log-uniform. So, the log-uniform prior is still used to set the time scale for all transitions leaving state *C*_2_. Then, we draw the statistical weights from the Dirichlet distribution for the second column of **K** such that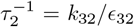. The other rates can be defined by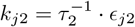. Whether we impose a log-uniform prior on 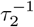 or some *k*_*j*2_ is irrelevant as long as this prior is introduced once for each column of **K**. We simulated the first binding rate (Model 1) with *k*_21_ = 2 · 600 *µ*M^*−*1^ s^*−*1^ which is the diffusion limit derived in [85] for ligand binding. The stoichiometric factor of 2 incorporates the structural information that two binding pockets are available. Of course the binding rate of a real process will be slower than the upper limit from [85]. To incorporate this aspect we take advantage of the fact that in our simulation of data and inference *k*_21_ = 2 · 600 *µ*M^*−*1^ s^*−*1^ is an arbitrary definition. In fact, we simulated *k*_21_ = 2 · 600 a. u. and defined this value to represent the value in SI units. We can also apply a different mapping of the arbitrary units to SI units. We may decide that 700 a. u. is supposed to represent the diffusion limit [85] in SI units. Thus we defined 700 a. u. = 600 *µ*M^*−*1^ s^*−*1^. In that way we avoid the unrealistic and extreme scenario that *k*_21,true_ is identical to the sampling box limit (which would still be for Bayesian inference a valid use case). All other “time-like” parameters such as sampling rates, dwell times and chemical rates need to rescaled by 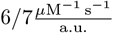.

The impact of the upper limits is shown in Fig. 3**a-b** and Fig. 4 (blue curves). The upper limits on the parameters *k*_21,max_ and *k*_32,max_ are now much smaller than the limits that we used previously. This solves the non-normalizability problem in the crucial parameter 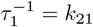 and reduces the chance that 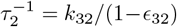 is non-normalizable. Still, 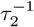 could diverge, if the marginal posterior of *ϵ*_32_ *→* 1 diverges. The parameters 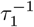 and *k*_32_ are now bounded from above and below by mathematically and physically motivated limits.

**FIG. 4.**
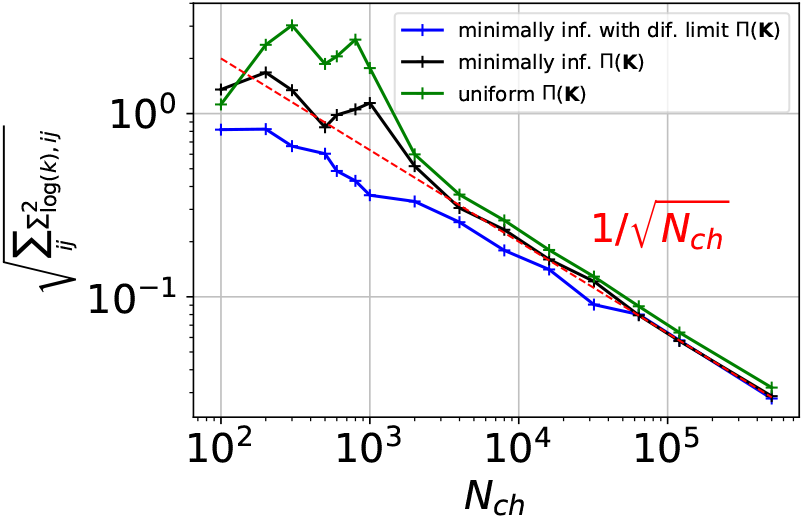
The uniform prior increases the posterior uncertainty. We define a multivariate analog of the standard deviation (Frobenius norm of the covariance matrix of the posterior) vs. *N*_ch_. The colors encode the prior assumptions. The larger the Frobenius norm, the more uncertainty remains after the inference. Plot is based on the same data used in Fig. 3.

The diffusion limits *k*_21_ *<* 2 · 700 a. u. and *k*_32_ *<* 700 a. u. restrict the RMSE (Fig. 3**a**, bluelines) to smaller values such that it drops below the 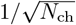 regime for *N*_ch_ *< N*_ch,crit_. However, even for data sets with *N*_ch_ *∈* [10^4^, 10^5^], the constraint adds information and decreases the ED. Constraining the two binding rates 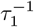 and *k*_32_ also influences via the likelihood the marginal posteriors of other parameters, particularly *p*(*ϵ*_21_ |*𝒴* _*T*_) (Fig. 3**b**). The constraints shape *p*(*ϵ*_21_ | *𝒴*_*T*_) into a more pronounced distribution which covers the true value.

#### 5. The uniform prior increases posterior uncertainty

The constraint only adds information to *p*(**K** | *𝒴*_*T*_) as long as the data *𝒴*_*T*_ themselves do not restrict the HDCV within the prior’s upper limits. To study the impact of the constraint it does not suffice to consider the RMSE. Notably, Error(log 𝔼 [***k***]) is not any quantity possible to report after an inference of real experimental data either. Ultimately, one judges the inference by its credibility intervals/regions (or confidence intervals/regions in an ML context) of the inferred parameters. Thus, only the posterior’s shape in general (the median/mean/peak, co-variance, and higher order statistical moments describing the tails) are at the modeler’s disposal to assess the quality of the inference.

Therefore, we define a quantitative measure of the spread of *p*(**K** | *𝒴*_*T*_), which is the Frobenius norm

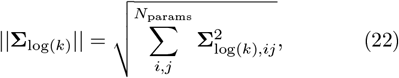

of the covariance matrix of the samples of *p*(**K**| *𝒴*_*T*_) on the log space of the chemical rates. The observed transition around *N*_ch,crit_ (Fig. 3**a**), that the location posterior derived from the uniform prior becomes equivalent to the posterior derived from Jeffreys prior without diffusion limit at *N*_ch_ = 2 · 10^3^, as judged by the RMSE, is only to some degree present in ||**Σ**_log(*k*)_ || (Fig. 4, black and green lines), which as noted measures the posterior’s spread not location. The spread of *p*(**K**| *𝒴*_*T*_) based on the uniform prior (green curve) does not converge to the spread of the minimally informative prior (black curve). It shrinks instead parallel given the *N*_ch_ range that is investigated. This effect becomes more prominent for the more complex CRN2. Further, the larger the size of the sampling box the larger ||**Σ**_log(*k*)_ || due to the practical non-identifiability problem. The Frobenius norm of the covariance of *p*(**K**| *𝒴*_*T*_) based on the Jeffreys prior, including the diffusion limit (blue curve), has prolonged smaller values than without the diffusion limit but finally converges towards the Frobenius norm of *p*(**K**| *𝒴*_*T*_) derived from the prior without upper limits (black curve). Hence, the information from the added diffusion limit is made use of, even up to *N*_ch_ = 3 · 10^4^. Above *N*_ch,crit_ the RMSE (Fig. 3**a**) is still a much more fluctuating parameter to benchmark the behavior of *p*(**K** | *𝒴*_*T*_) than the rather non-fluctuating ||**Σ**_log(*k*)_||. ||**Σ**_log(*k*_||_)_ follows almost a straight 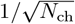-scaling (Fig. 4). In other words, repeating the experiment under identical conditions will leave ||**Σ**_log(*k*)_ || almost unchanged, but the posterior’s location will be randomly translocated from data set to data set within the soft constraints of **Σ**_log(*k*)_. Larger simulation boxes increase the value of the green curve the most, followed by a minor effect on the black curve. This is an elegant result because, ultimately, with real experimental data, the inference quality is judged by uncertainty quantification and not by the RMSE.

### D. The minimally informative prior still generates an improper posterior making it difficult to decide which point estimate has the smallest error

To confirm that *p*(**K**| *𝒴*_*T*_) is still improper (Eq. 8) and where in the parameter space *p*(**K**) dominates *p*(**K**| *𝒴*_*T*_), the most worrying 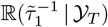 and ℝ (*k*_32_|*𝒴* _*T*_) are plotted for *N*_ch_ *<* 2 · 10^3^ and sampled in larger sampling boxes (Fig. 5). The PC data originates from the 4-states-1-open-state CRN (Fig. 2 **a**). Note that 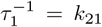 because only one Markov transition leaves state *C*_1_.

**FIG. 5.**
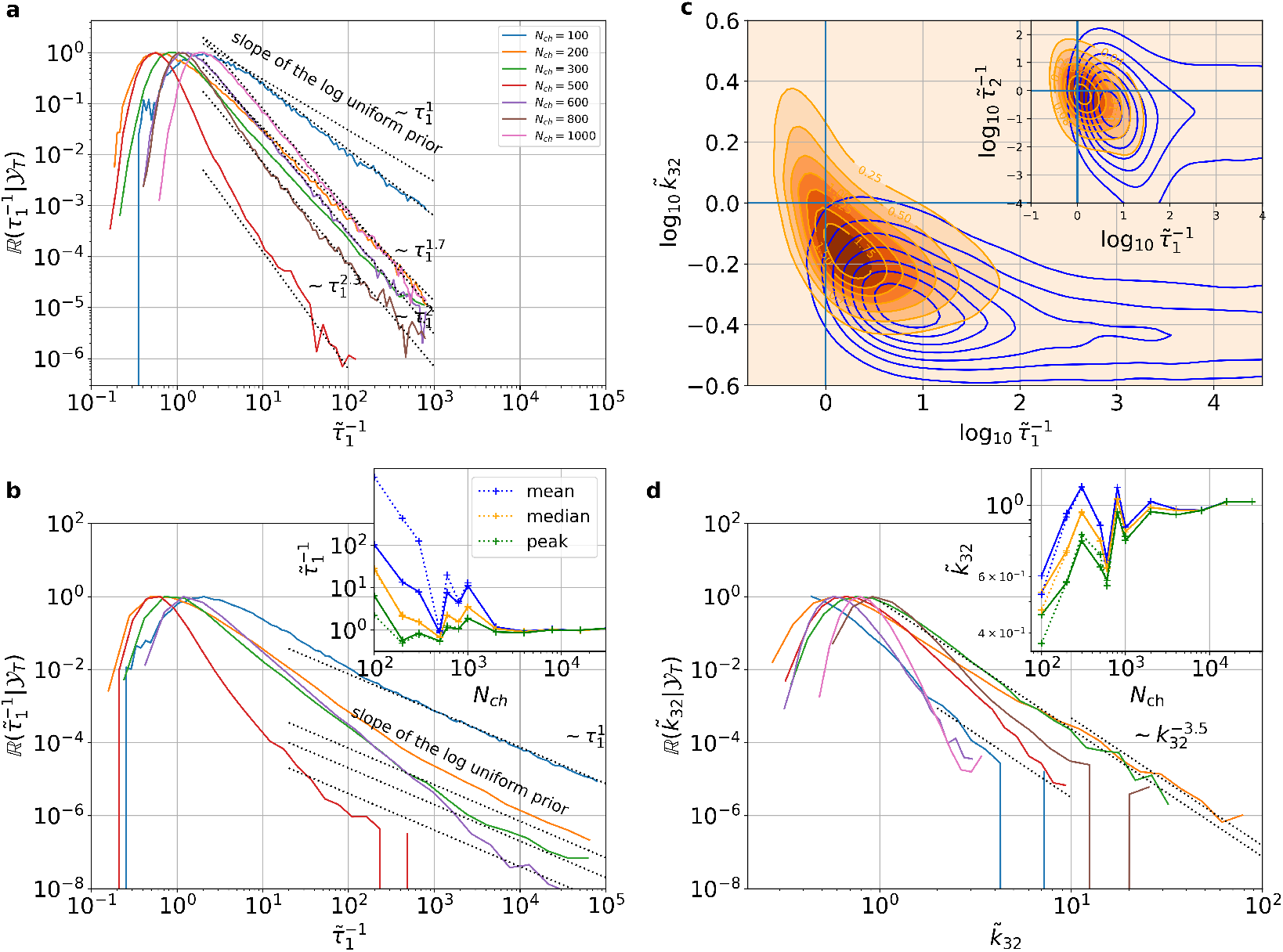
The minimally informative prior without diffusion limit generates still improper posteriors for CRN1. The relative marginal posteriors 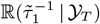 and 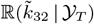 are plotted for different *N*_ch_ and different sampling boxes. The tilde over the parameters indicates that each parameter is normalized to its true value. **a** 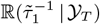displays power law-like behavior, but appears proper given the limits of the sampling box. **b**, However, the same 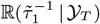sampled from a two magnitudes larger sampling box displays an area for large 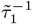 where the prior entirely dominates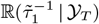. The inset shows the mean, median, and peak for 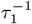vs. *N*_ch_ for the smaller sampling box (solid line) and larger sampling box (dotted) line. **c**, 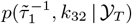for *N*_ch_ *∈{* 100, 200*}* (blue, red densities) displays a non-local correlation structure leading to a bias of ℝ (*k*_32_| *𝒴*_*T*_). The inset is based on the same data *N*_ch_ *∈ {*100, 200*}* but *k*_32_ is transformed to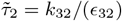. **d**, For ℝ (*k*_32_|*𝒴*_*T*_) the heavy tails are much less a concern. The power law exponent for the smallest *N*_ch_ is *ϕ ≈* 3.5. The insets of the panels display the mean (blue), the median (orange), and the peak (green) of the marginal posterior.

As a guide to the eye, functions *f* (*τ*^*−*1^) *∝* (*τ*^*−*1^)^*−ϕ*^ (dashed lines, Fig. 5**a-b**,**d**) that have a power law scaling are plotted, including the log-uniform prior. The right tails of 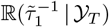 are well approximated by different power laws until the limit, 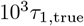, of the sampling box (Fig. 5**a**). The slopes of the tails are heavily influenced by the log uniform prior 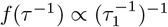 but also the likelihood contributes some additional slope (in other words information) till approximately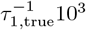. For the smallest sampling box, for all values of *N*_ch_ the 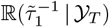 seem to be proper but only vaguely identified. The exponent *ϕ* gradually increases with *N*_ch_ indicating the increase of information about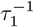. However, using a two-orders of-magnitude larger sampling box (Fig. 5**b**) demonstrates that for even larger 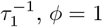holds. I.e. the prior complete dominates *p*(**K**|*𝒴* _*T*_) in this region of **Θ** in the direction of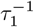. Hence, *p*(**K**|*𝒴*_*T*_) indicates a practical non-identifiability problem of the HMM as only the prior (Eq. 10) and not the likelihood contributes to the posterior locally. For at least *N*_ch_ *≤* 600 the data do not contribute information for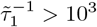. The limited influence of some chemical rates or dwell times in some parts of **Θ** on the signal is a general feature of partially observed CRNs (App. B). In the inset of Fig. 5**b**, the mean (blue), the median (orange), and the peak (green) of 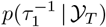vs. *N*_ch_ for the smaller sampling box (solid) and larger simulation boxes (dashed) are plotted. Mean values or higher statistical moments do not exist for distributions with power law tails with exponents *ϕ <* 2, thus the peak should be reported for the barely identified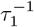. Consistently, the peak of 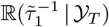 (green curve inset Fig. 5 **b**) has maximally a relative error of a factor of 2 and seems unbiased and not diverging with increasing sampling box size, given the tested data and minimally informative prior. For the same data, the residuals of the mean (blue curve inset Fig. 5 **b**) of 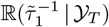 are heavily biased, sensitive to the sampling box limits and orders of magnitude away from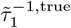. The median (orange curve inset Fig. 5 **b**) is more robust against different sampling box sizes. However, it is still biased towards too large values of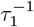, which is unsurprising as it is not defined on the full support (0, *∞*) if the shape of the log uniform prior [86] eventually dominates 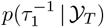 for large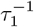. This potentially indicates for parameters with strong practical non-identifiability degree (the magnitude of the posterior peak does not decay by multiple decades before reaching the area where the prior dominates the posterior) that it is better to report MAP values. In Fig. 5**c** we plot 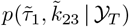 and in the inset 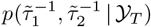to demonstrate that the practical non-identifiability of 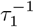 leads to bias of *k*_32_ however the bias is much reduced for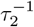. The bias of the corresponding *p*(*k*_32_| *𝒴*_*T*_) is also present for different data sets (inset of Fig. 5**d**).

The tails of ℝ (*k*_32_|*𝒴* _*T*_) given the sampling box, seem to be less heavy tailed (Fig. 5**d**) than those of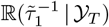. For *N*_ch_ =*∈ {*100, 200, 300*}*, R(*k*_32_ | *𝒴*_*T*_) appears to follow a power law, hence, 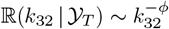 with *ϕ* = 3.5. The quicker decaying power-law tails of R(*k*_32_| *𝒴*_*T*_) still create a skewed distribution. The skewed tail of R(*k*_32_|*𝒴* _*T*_) compensates partially for the bias of the peak region of ℝ (*k*_32_|*𝒴* _*T*_) (Fig. 5**d**). Hence, 𝔼 [*k*_32_] (blue curve, inset, Fig. 5**d**) is the least biased point estimator towards too small *k*_32_ values until the data are strong enough. In contrast to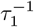, the standard Bayesian point estimate, 𝔼 [*k*_32_] performs best for *k*_32_. Besides our observation, mathematics dictates that reporting *θ*_MAP_ for parameters whose posterior has a powerlaw tail with small *ϕ ≤* 2 is more robust because mean values and eventually medians for *ϕ ≤* 1 do not exist. However, it is unclear whether reporting the mean value for parameters with large *ϕ* because of their smallest bias (inset of Fig. 5**d**). generalizes to other CRNs. Note that from the bias of the peak of ℝ(*k*_32_|*𝒴*_*T*_) to too small values, it is clear that small HDCIs around the peak will not include the **K**_true_ with the frequency equal to the probability mass they claim to have. Thus, posteriors which fullfill decently the asymptotic behaviour of the Berstein-von-Mises theorem need larger *N*_ch_ or a second observable [38].

In summary, the minimally informative prior, particularly, its convex log uniform distribution for *τ*_*i*_’s or *k*_*ij*_’s has the desirable feature of concentrating *p*(**K**|*𝒴*_*T*_) much closer to **K**_true_, but it still produces improper *p*(**K**| *𝒴*_*T*_) if no further upper limits can be justified. The minimally informative prior alleviates the improperness of *p*(**K** | *𝒴*_*T*_) by making the posterior less sensitive to the often nonphysical and arbitrary limits of the sampling box, but the practical non-identifiability problem will become relevant when increasing the sampling boxes at some point for all data sets. Further, the higher the data quality, the less sensitive is the inference to the sampling boxes limits. Hence, the degree of the practical non-identifiability problem has to be judged based on how much the peak of 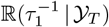 decays before the slope of 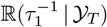 is essentially the slope of the prior.

### E. Solving the practical non-identifiability problem with vague additional information on cooperativity

We exemplify how to robustify an improper *p*(**K**| *𝒴*_*T*_) by physically justified vaguely informative prior distributions. For CRN1, we enforce a soft physical constraint on the one hand on the binding rates *k*_21_ and *k*_32_ and on the other hand on the unbinding rates *k*_12_ and *k*_23_ by a regularizing prior, plausible for homomeric proteins. Physical common sense dictates that one should be skeptical *a priori*, if binding rates or unbinding rates for the same/similar binding sites have values differing by orders of magnitude. One modeling assumption encoding this skepticism could be

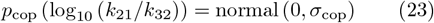

for binding and for unbinding

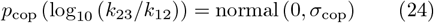

with some finite standard deviation *σ*_cop_ (Fig. 6). Note that this corresponds to the classical definition of cooperativity: The ratio of *K*_32_ to *K*_21_: If the affinity *K*_*ij*_ = *k*_*ij*_*/k*_*ji*_ increases with the number of occupied binding sites, this is called positive cooperativity. I.e. if the microscopic binding rates(the binding rates per binding site) are constant, e.g. diffusion limited, the ratio *r* = *K*_32_*/K*_21_ = *k*_12_*/k*_23_ can be used as measure of cooperativity with = 1 equals no cooperativity, *>* 1 is positive, *<* 1 negative cooperativity. The vaguely informative prior is a much less radical prior (assumption) than assuming identical microscopic rates (which is the non-cooperative assumption, *σ*_cop_ = 0, for the binding and unbinding), as frequently done for ligand gated [87, 88] and more excessively for voltage gated ion channels [89–92] to alleviate structurally non-identifiable or practical non-identifiability problems in HMM inferences. The CRN of the shaker channel [90] is a good example of how the structural prior information of having four subunits within the shaker channel implies a necessary complexity of the CRN to allow it to explain the data and to be physically interpretable. This CRN has that many voltage dependent rates that setting certain subsets of rates equal is used to avoid non-identifiable problems. However, this assumption might be incorrect for the ion channel at hand. Thus one gains identifiably of the parameters by loosing potentially the ability of the model to express the true process. Instead, assuming that cooperativity cannot change the chemical rates beyond some reasonable scale is physically plausible and restricts the model much less. One might debate the prior’s variance and the prior’s shape. Note that adding this additional regularizing prior solves the improperness problem of 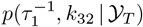originating from a flat likelihood for high values of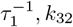 no matter what finite *σ*_cop_ is used because of the quickly decaying tails of normal distributions. Notably, almost any additional prior that adds at least a tiny amount of decay till infinity to the otherwise log uniform dominated posterior would also render *p*(**K**|*𝒴*_*T*_) proper. The normal distribution on the logspace has, by definition, desirable properties: It is symmetrical on the order of magnitude, and defines an area, within one standard deviation, with little impact on *p*(**K**|*𝒴*_*T*_) and an area of increasing penalty for values away by multiple standard deviations. In that way extreme conclusions from the data, expressing strong cooperativity effects in the data-generating process, have to be more and more supported by the data to be represented with a relevant magnitude in *p*(**K**| *𝒴*_*T*_). Using the larger sampling box (Fig. 6) makes the regularization more urgent. In Fig. 6 panel a,b, we demonstrate how applying the additional prior renders 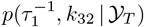 proper even for *N*_ch_ = 10^2^ and *σ*_cop_ = 1. Notably, in this case *± σ*_cop_ covers 2 oders of magitude. The bias (inset, Fig, 6**a**) of *p*(*k*_32_| *𝒴*_*T*_) is drastically reduced with a finite *σ* compared to the pure minimally informative prior, and decreasing *σ* further improves the inference (Fig. 6**b**) in terms of bias and variance. Notably, the prior nudges the posterior to concentrate its mass between 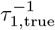 and *k*_32,true_. Thus, at some point, the smaller *σ*_cop_ of Eq. 23 and 24, the more is the variance of the posterior decreased but also the bias of *p*(log_10_ (*k*_23_*/k*_12_)) starts to dominate the posterior more and more. The traditional non-cooperativity assumption trades variance of *p*(**K**|*𝒴*_*T*_) for a maximum of bias and thus should only be applied if there is strong *a priori* evidence that non-cooperativity is true in the data-generating process.

**FIG. 6.**
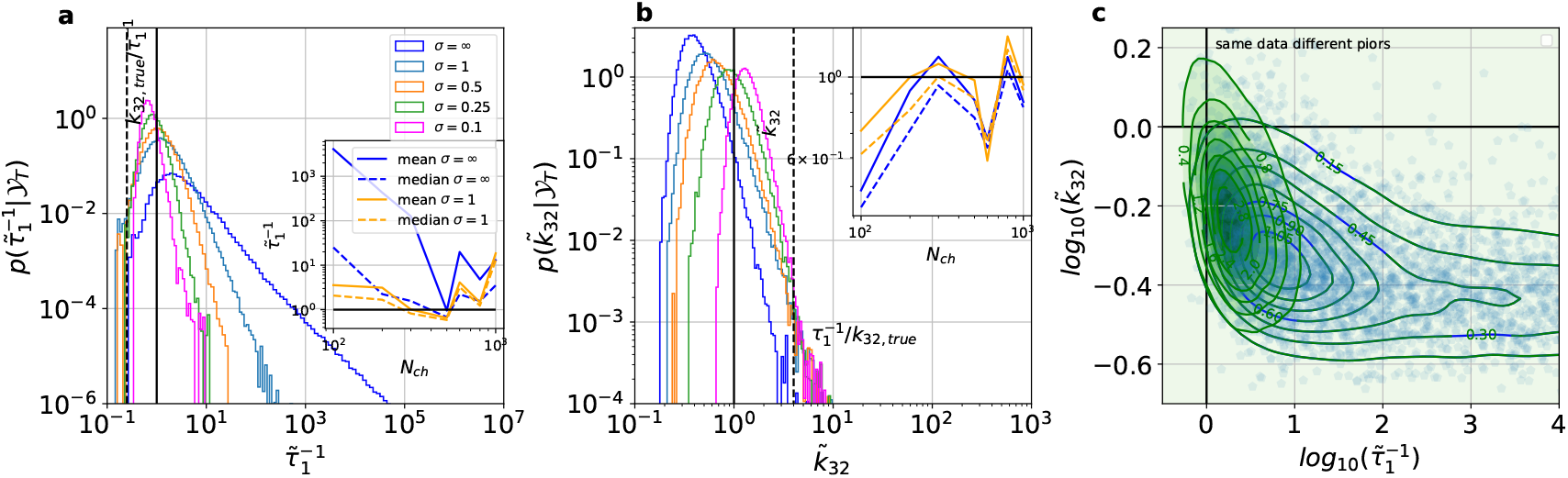
A vaguely regularizing prior on the cooperativity factor renders the posterior proper even for the lowest quality data. We demonstrate for the previously used data sets of CRN1 (Fig. 2**a**) with *N*_ch_ = 10^2^ the effects of the value of *σ*_cop_ using the larger sampling box. Note that 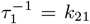 holds. The black continuous lines indicate the value of 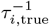 and *k*_32,true_ on the x-axis. The black dashed lines indicate the corresponding 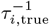 and *k*_32,true_ in units of the true value of x-coordinate to visualize the bias of the regularizing prior, constraining the posterior more and more between the dashed and solid black line with decreasing *σ*_cop_. **a**, 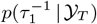for either no or a series of increasingly strong regularization. The inset compares the effect of the vague regularization (orange) on the mean (solid curve) and median (dashed curve) vs. *N*_ch_ with no regularization (blue). **b**, *p*(*k*_32_|*𝒴* _*T*_) for the same series of increasing regularization. The inset compares the effect of the vague regularization (orange) on the mean (solid curve) and median (dashed curve) vs. *N*_ch_ with no regularization (blue). **c**, The effect of the regularization on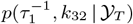, with no regularization (blue) and the most vague regularization prior (*σ*= 1, green).

### F. Minimal informative prior to ward off the curse of complexity and dimensionality

Next, we investigate the challenges arising when the complexity of **K** is increased. CRN2 additionally contains two flip states (Fig. 2 **d**). We enforce microscopic reversibility, which reduces the number of to-be-inferred parameters by one. For a comment on how the minimally informative prior improves the convergence of the sampler to the true posterior (Ap. F 1) and alleviates *the curse of dimensionality*.In Fig. 7 **a** the pathological dimensions of *p*(**K**| *𝒴*_*T*_) derived from the uniform *p*(**K**) (green) against the same dimensions of *p*(**K**|*𝒴* _*T*_) (black) derived from the minimally informative *p*(**K**) are compared. The less pathological dimensions, in the sense that they deliver roughly Gaussian marginal posteriors, are plotted in Fig. 7 **b1-6**. Even for *N*_ch_ = 10^5^, the posterior based on the uniform prior clearly demonstrates the practical non-identifiability feature of the likelihood in 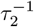 and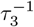. The ridge-like correlation of 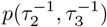(panel **a3**) with a slow decay along the ridge demonstrates the practical non-identifiability problem. Notably, the ridge is strongly constraint for values smaller than *τ*_2,true_ and *τ*_2,true_ (panel **a1-2**) but above it seems to extend to infinity. The strong correlation between 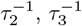 produces a paradoxical challenge of the minimally informative prior which turns out problematic for smaller *N*_ch_. The result of the strong correlation is that the sample value for 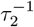 can be predicted by a affine linear function 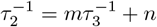 (panel **a3** dashed line). Ignoring the affine part of the function, the slope of loguniform prior on 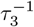 will be mapped by the linear function to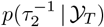. Additionally, 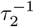 has its own log uniform prior. The same applies to both parameters the other way around. Hence, both one-dimensional priors together with the practical non-identifiability ridge, contribute to the posterior a scaling of 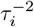 for both parameters, creating bias towards too small values (see, the corresponding 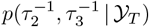 in Fig. 8 **a**). Based on the corresponding 2d marginal distributions (Fig. 7 **a**), the Markov transition probabilities *ϵ*_12_, *ϵ*_32_ and *ϵ*_23_ appear to have a more and more diverging posterior the larger the sampling box for 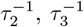 gets. These parameters should be considered unidentified, based on the herein-employed heuristic and visual criterion (see discussion around Eq. 9) to asses the degree of practical non-identifiability *p*(**K**|*𝒴* _*T*_). Surprisingly, 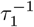 is for CRN2 not as unidentified as for CRN1 (Fig. 3 **b**). We employed a smaller sampling box for the uniform prior (because of an often not converging sampler, likely caused by the practical non-identifiability problem and the curse of dimensionality.Using a smaller sampling box for the uniform prior also disadvantages the minimally informative prior in comparison because less of the part of the parameter space where the likelihood is flat is possible for the sampler to reach. In the App. Fig. 15 we demonstrate that *sufficiently proper* posteriors are achieved, for the minimally informative prior at the experimentally possible, but certainly challenging data set of *N*_ch_ = 5 · 10^3^. However, with the uniform prior one needs an impossible data quality of *N*_ch_ = 2.5 · 10^5^. Hence, the minimally informative *p*(**K**) increases the range of acceptable data for this CRN roughly a 50-fold. Using only the minimally informative prior (black posterior) produces for *k*_32_ and *k*_45_ visually-visible improper marginal posteriors for *N*_ch_ *<* 5 · 10^3^ (see Fig. 7 and 8).

**FIG. 7.**
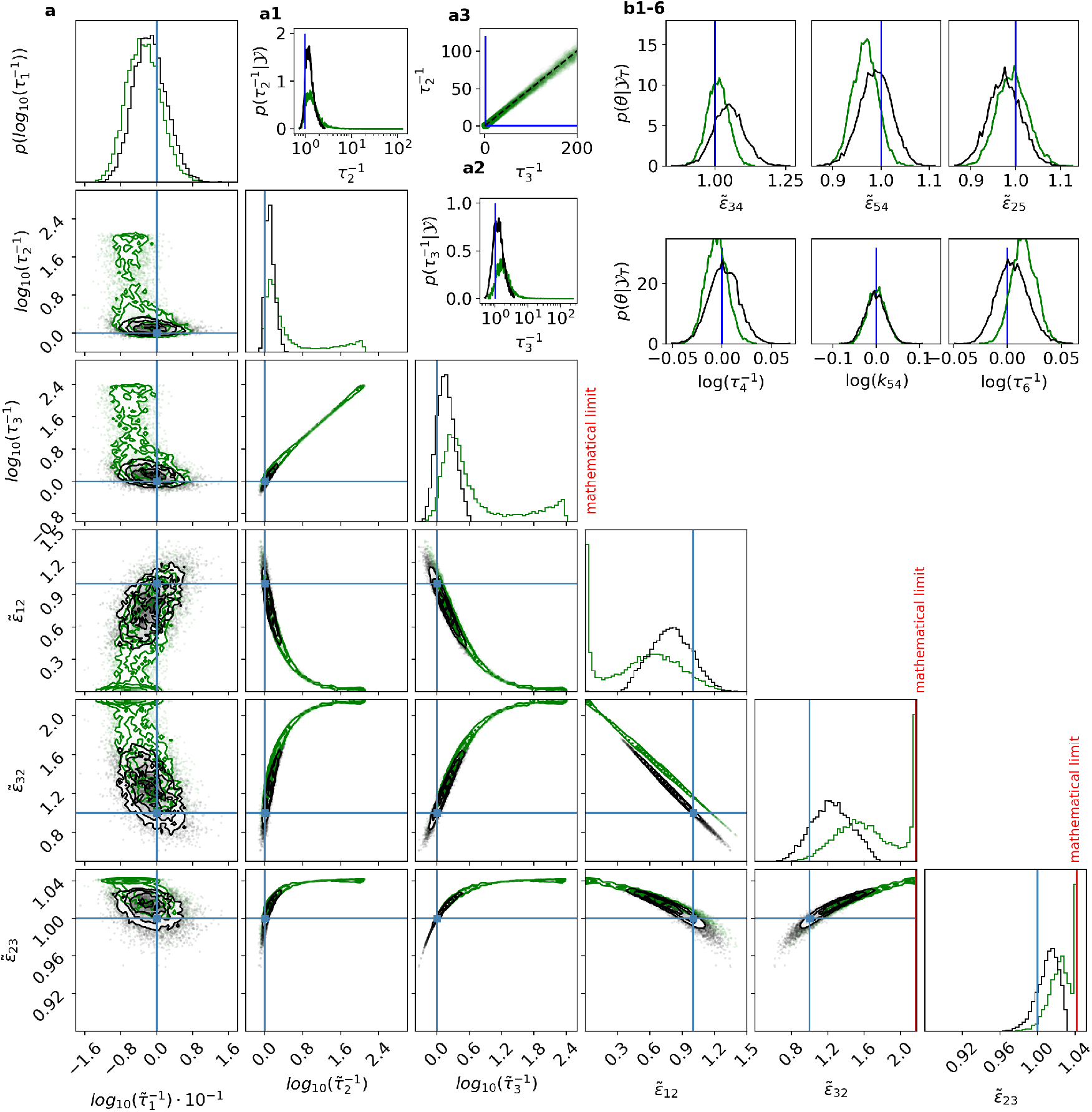
For the more complex CRN2 the minimally informative prior is necessary even for data sets of of unrealistic high quality such as *N*_ch_ = 10^5^. **a**, Posteriors based on uniform (green) and minimally informative priors (black) are compared. The clearly non-Gaussian-shaped marginal posteriors plus those concerning 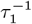 are plotted in **a**, all other, rather Gaussian marginal posteriors, are shown in **b**. The insets **a1, a2** visualize the posterior without the log transformation. The black posterior is equipped with the minimally informative *p*(**K**). The green posterior is based on the uniform *p*(**K**). The flat (green) posterior for 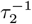 and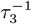, creates what appears to be a exponential increase for 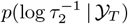 and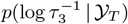. **a3**, demonstrates the positive linear correlation contained in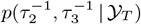. Deviations of both parameters from the corresponding true value can be compensated to some extent by the other parameter. Note that the sampling box for the posterior samples of 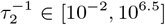 updated from the minimal information is more than an order of magnitude larger than the range of the simulation box used when the uniform *p*(**K**) is used. This disadvantages the posterior based on the minimally informative prior but demonstrates the larger robustness of the minimally informative prior. The posteriors derived from the uniform *p*(**K**) are sampled by *k*_*ij*_ and then mapped to the (***τ***, ***ϵ***)*−*space.

**FIG. 8.**
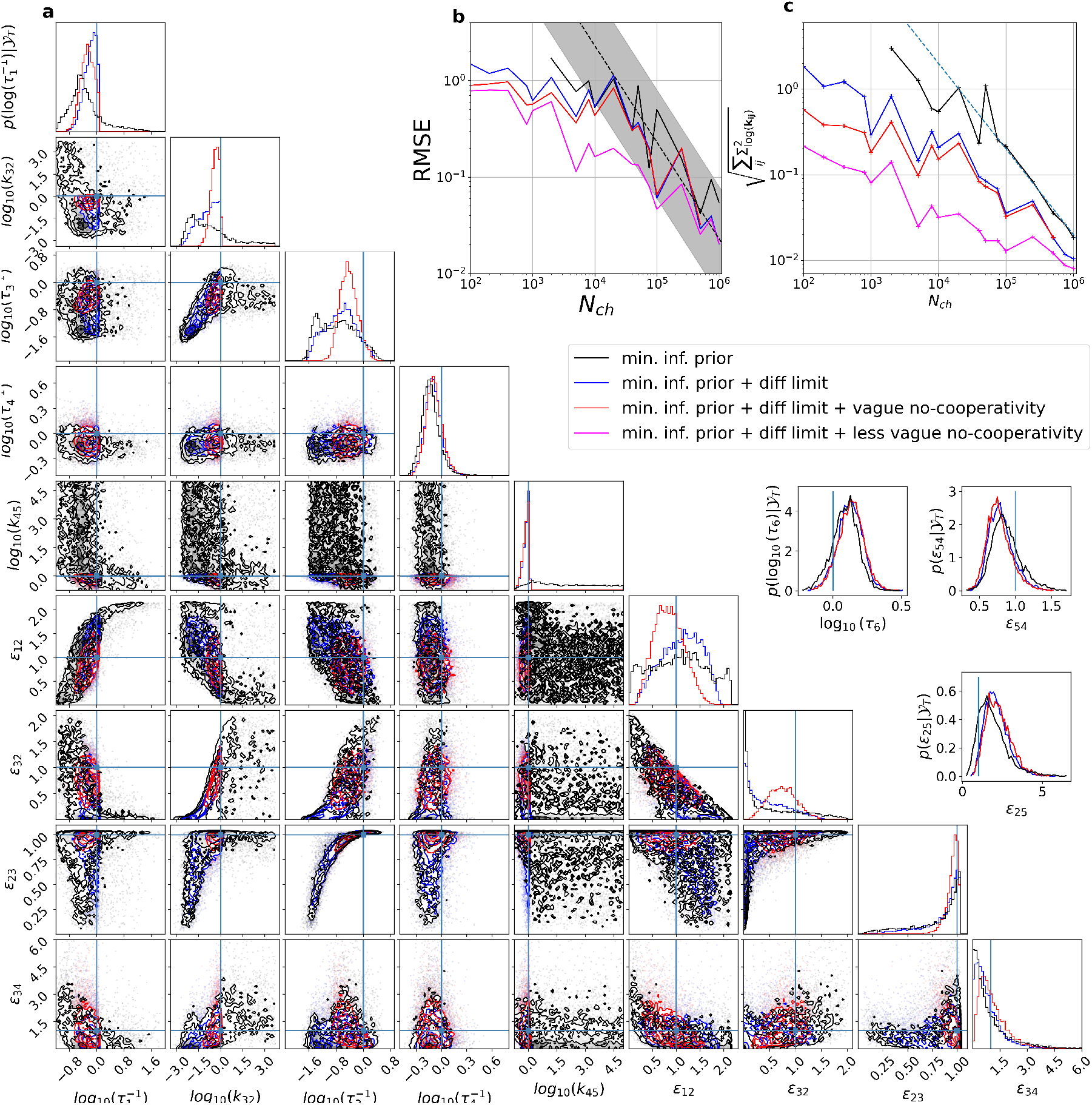
For CRN2, the minimally informative prior enables inference for about two orders of magnitude lower data quality. Adding information by diffusion limits and vague bias towards non-cooperativity allows us to work with three orders of magnitude lower data quality. The data sets simulated from CRN2 (Fig. 2) are analyzed The color black refers in all plots to *p*(**K**| *𝒴*_*T*_) based on the minimal informative prior. Blue corresponds to assuming diffusion-limited binding. Red an additional assumed vague prior on the cooperativity of the binding and unbinding rates. Magenta less vague no-cooperativity assumption. **a**, The true values of **K** are indicated by the blue lines. Posterior for *N*_ch_ = 10^2^ for the minimally informative prior, minimally informative with upper limits and with an added vague no-cooperativity assumption. For visual clarity, we suppress 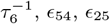 in the main plot but add sub panels which display the corresponding posteriors. Note that these parameters are only slightly influenced by the priors, and even without the priors, the posterior is peaking Gaussian-like with some skewness. **b**, The RMSE of the log space of the chemical rates is plotted vs. *N*_ch_ for the median (solid curve) and the marginal peak (dashed curve) for different prior assumptions. **c**, The Frobenius norm of all *k*_*ij*_ s of the covariance matrix of the samples of **K**

### G. Effects of combining the theoretical diffusion limit with vague non-cooperativity assumptions

In Fig. 8 **a** we display *p*(**K** | *𝒴*_*T*_) for the smallest tested data quality *N*_ch_ = 10^2^ with the minimally informative prior and with the additional (vague and hard) constrains discussed below. Using only the minimally informative prior (black posterior) produces for *k*_32_ and *k*_45_ not sufficiently proper marginal posteriors. The first *visually* proper *p*(**K** |*𝒴*_*T*_) appears with *N*_ch_ = 5 · 10^3^ (App. Fig. 15 **a**). From the discussion before, it is clear that the likelihood for *N*_ch_ = 5 · 10^3^ is only *sufficiently* practical identifiable that the improperness of *p*(**K**| *𝒴*_*T*_) is not detected visually even though there is no reason to assume that it is not there if one would sample from much larger simulation boxes. Due to the complexity of the inference problem we employ the additional assumptions.

#### 1. The vague no-cooperativity assumption increases the accuracy and decreases the uncertainty of the inference

If one applies strict upper diffusion limits for *k*_21_, *k*_32_ and *k*_45_ (Fig. 8, blue posterior and curves) one gains a proper *p*(**K**| *𝒴* _*T*_) for *N*_ch_ = 10^2^ in the corresponding dimensions and also the other dimensions become *sufficiently proper*. Adding further vague prior assumptions log_10_(*k*_21_*/k*_32_) ∼ normal(0, 0.5), log_10_(*k*_12_*/k*_23_) ∼ normal(0, 0.5), log_10_(*k*_45_*/k*_32_) ∼ cauchy(0, 1) and log_10_(*k*_54_*/k*_23_) ∼ cauchy(0, 1), which regularizes *p*(**K**| *𝒴*_*T*_) gently towards a non-cooperative CRN (Fig 8 red posterior), reduces the mean-distance between *p*(**K**| *𝒴*_*T*_) and **K**_true_ and reduces the uncertainty (Fig. 8 **c**). Note, that a less-vague no-cooperativity assumption log_10_(*k*_21_*/k*_32_) ∼ normal(0, 0.1), log_10_(*k*_12_*/k*_23_) ∼ normal(0, 0.1) reduces the Frobenius norm and RMSE further (see 8 **b-c**).The RMSE (Fig. 8 **b**) shows that at *N*_ch_ *≈* 5 · 10^4^ the RMSE transitions to 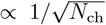 asymptote while for*N*_ch_ *<* 5 · 10^4^ the practical non-identifiability problem combined with prior assumptions influence the posterior. Above *N*_ch_ = 5 · 10^4^, the posteriors equipped with priors with diffusion limit produce similar RMSEs, unless the less vague cooperativity assumptions (Fig. 8 **b** magenta curves and posterior) are used. For the less vague prior the RMSE converges onto the curves of the other two prior assumptions (blue and red curve) around *N*_ch_ = 10^6^. In contrast, the pure minimally informative prior has different RMSEs (Fig. 8 **b** black curve) for each data set. This shows that the vague no-cooperativity assumptions lost their influence on the RMSE, while the diffusion still influences the RMSE.

The Frobenius norm of the covariance matrix of *p*(**K**|*𝒴* _*T*_) shows (Fig. 8 **c**) that enforced upper diffusion limits (blue, red, and magenta curve) still add information and reduce the uncertainty of *p*(**K**| *𝒴* _*T*_). Hence, even for data qualities of *N*_ch_ *>* 10^6^, an ML inference would ignore relevant information to reduce the uncertainty of the inference. The Frobenius norm of the posteriors based on the pure minimally informative prior without additional assumptions transitions at *N*_ch_ = 10^5^ to the 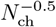-scaling.

To summarize, with minimally informative prior with diffusion limits (Fig. 8), one can make inferences with more than 10^3^ times smaller *N*_ch_ per time trace compared to Bayesian inferences with the uniform prior (Fig. 7) or ML/MAP inferences. Note that theoretical data qualities of *N*_ch_ *>* 10^4^ are beyond experimentally achievable data qualities. The added vague non-cooperativity prior contribute information to the posterior approximately until *N*_ch_ = 10^5^ as judged by the RMSE and Frobenius norm. For the less vague prior the gain of relevant information lasts for values even higher than *N*_ch_ = 10^6^ as judged by the Frobenius norm, but the RMSE are roughly the same.

#### 2. To what extent can channel properties be assessed against the bias of the no-cooperativity prior?

We test for the different no-cooperativity priors, what typical data quantity is needed such that *p*(**K** |*𝒴* _*T*_) supports positive cooperativity (defined as *r* = *k*_12,true_*/*(*k*_54,true_) *>* 1 Sec. V E). One could also ask at what point becomes the bias of the additional soft coupling by the prior towards no-cooperativity detrimental to the inference because the DGP (Fig. 2) is cooperative with *k*_12,true_*/k*_54,true_ = 5*/*2. There are essentially two categories (*negative k*_12_*/k*_54_ *<* 1 and *positive k*_12_*/k*_54_ *>* 1) and the infinitesimal thin (green) line in between with no *cooperativity k*_12_*/k*_54_ = 1. Let *r* = *k*_12_*/k*_54_ be our measure of coorperativity. If

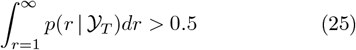

the data supports positive cooperativity Notably, if 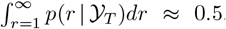, also a negative cooperativity model is plausible just as positive cooperativity. Following the Bayesian paradigm, we are not looking for any binary significant test result, but embrace the continuous aspect of the question at hand. The inequality 25 is fullfilled if the posterior median (solid lines) holds med_*post*_(*r*) *>* 1. We contrast the median with HDCIs that tell *what is the smallest most probable interval*. For skewed posteriors (because of uninformative data), HDCIs might indicate a different cooperativity model than the median. Note that for small *N*_ch_ we work in a regime where frequentist testing would likely not produce significant results.

For the pure minimally informative prior (Fig. 9 **a**), and *N*_ch_ *∈* [5 · 10^3^, 2 · 10^4^] the posterior flucuates between a weak indication based on the median (solid line) that there is positive cooperativity and that both models are equally plausible. In particular, the median is biased towards too small values of *r*. The 0.5-HDCI (lower limit dashed black lines) is almost entirely smaller than *r* = 1. To have a unbiased median and a 0.5-HDCI that additionally supports qualitatively positive cooperativity one needs at least *N*_ch_ *>* 5 · 10^4^. In contrast, working with the uniform requires *N*_ch_ = 2.5 10^5^.

**FIG. 9.**
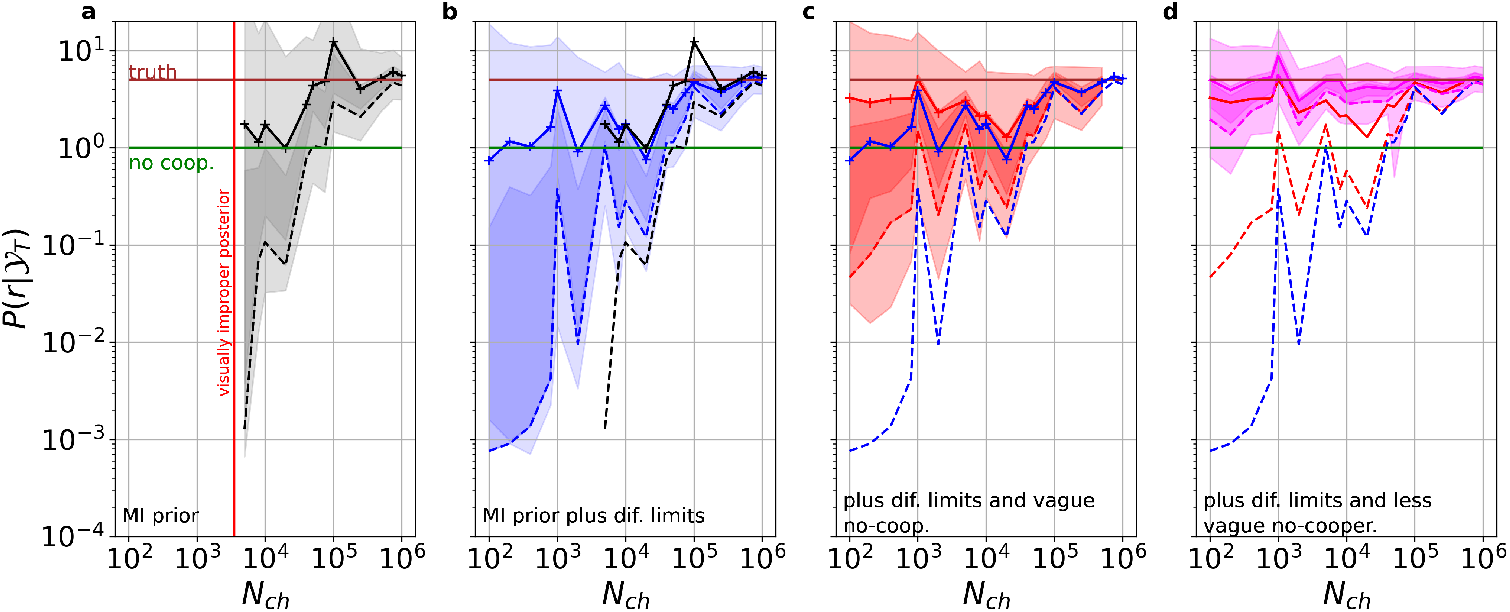
The prediction of positive *cooperativity* (accelerated unbinding) of the unbinding of the second ligand gains certainty with vaguely informative prior assumptions with a bias towards no-cooperativity. On the x-axis we plot *N*_ch_ as a proxy for the information content in the data originating from CRN2. The dashed lines indicate the lower limit of the 0.5-HDCI, and the solid lines are the medians. The color corresponds to the prior assumptions with more information added from left to right. The set of three [0.99,0.5,0.3]-HDCIs is plotted in each panel. As a guide to the eye to discern the gain of certainty that there is negative cooperativity and the reduction of bias, we replot the median and lower limit of the 0.5-HDCI from the previous panel. For visual clearity we supress the black lines in panel **d. a**, minimally informative prior. **b** With an additional diffusion limit assumption. **c** With additional vague no-cooperativity assumption log_10_(*k*_21_*/k*_32_) ∼ normal(0, 0.5), log_10_(*k*_12_*/k*_23_) ∼ normal(0, 0.5), log_10_(*k*_45_*/k*_32_) ∼ cauchy(0, 1) and log_10_(*k*_54_*/k*_23_) ∼ cauchy(0, 1). **d** With an additional less vague no-cooperativity assumption log_10_(*k*_21_*/k*_32_) ∼ normal(0, 0.1), log_10_(*k*_12_*/k*_23_) ∼ normal(0, 0.1). A standard deviation of these priors corresponds approximately to 3.1 relative deviation of the corresponding parameters. Further, log_10_(*k*_45_*/k*_32_) ∼ cauchy(0, 1) and log_10_(*k*_54_*/k*_23_) ∼ cauchy(0, 1).

One may ask where the bias towards too small *r* values for *N*_ch_ *∈* [10^3^, 4 · 10^4^] comes from. The normal approximations used to derive the likelihood [38] is justified given the scale of *N*_ch_. Thus, we suspect the bias to originate from correlations and a strong practical non-identifiability problem in the likelihood/posterior. Imagine a strong positive correlation after the inference between, e.g., *τ*_1_ and *τ*_2_ such that one can predict very accurately from *τ*_1_ the scale of *τ*_2_. The limiting extreme case of such a correlation between two parameters is realized in the structurally non-identifiable problem of the linear birth-death model (Eq. 1). Giving *k*_birth_ a log-uniform prior would result in *k*_death_ having a log-uniform prior. Supplying to both parameters log-uniform priors results – after the inference – in an effect of the combined prior on the posterior as 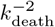 and equivalently 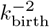. An anticorrelation between the parameters would eliminate the effect of the prior. Adding the diffusion limit to the posterior (Fig. 9**b**) extends the region of a visually *sufficiently proper* posterior (Fig. 8) at least to *N*_ch_ = 10^2^ and decreases uncertainty (difference between black and blue dashed line). But an effect of the capabilities of the median to predict qualitatively positive cooperativity is small or not existent. While not necessarily indicating a wrong model, the median is typically undecided and thus biased to too small *r* values. Hence, adding the diffusion limit reduces uncertainty of *p*(*r*|*𝒴* _*T*_) but does not help to answer the fine-grained question of cooperativity, unless one works with unrealistic data qualities (see uncertainty Fig. 9**b** above *N*_ch_ *>* 10^5^).

This changes when adding the vague no-cooperativity prior (Fig. 9 **c**), which by its definition biases *p*(*r*| *𝒴*_*T*_) more around the green line. The median and the HDCIs (see, Fig. 9 **c** the difference between red and blue lines) are shifted towards *r*_true_ against the bias of the Cauchy prior acting on *r*. The median now always indicates positive cooperativity. The 0.5-HDCI is, roughly speaking, undecided but much less biased without the vague regularization.

We show in Fig. 9**d** the effect of decreasing the variance of the prior for the ratios (increasing the bias towards non-cooperatively of the binding and unbinding rates) on the one hand between *k*_12_ and *k*_23_ and on the other hand between *k*_21_ and *k*_32_, while the cooperativity priors for o*k*_45_ and *k*_32_ and *k*_54_ and *k*_23_, remain the same. This structure vaguely incorporates the prior knowledge that the vertical Markov transitions (Fig. 2 **c**) in the CRN represent changes in the protein, which might alter the binding and unbinding rates to some amount. A further shift of the lower limit of the 0.5-HDCI (dashed lines) and the median (dotted lines) can be observed. For realistic PC data quality assumptions *N*_ch_ *<* 10^4^ the prior combining diffusion limits with less vague non-cooperativity assumptions performs the strongest but also supposes the most. Only for high and unrealistic data quality *N*_ch_ *>* 10^5^ the posteriors without the vague non-cooperativity assumptions (black) seem to have a smaller bias to smaller negative cooperativity.

## VI. CONCLUSION

Bayesian inference offers efficacious remedies for practical non-identifiability problems in HMM inference thereby allowing parameter uncertainty quantification for finite data. Nevertheless, pathologies of the likelihood also pose challenges for Bayesian inference.

If little about the actual values of some parameters is known *a priori*, we show that minimally informative priors are crucial to expand the range of acceptable data quality. They attempt to make posteriors as sensitive to the data as possible, thereby also alleviating practical non-identifiability pathologies in HMMs. The suggested minimally informative prior increases accuracy and decreases uncertainty compared to a uniform prior.

Any prior dominates the posterior in the regions of a constant likelihood value (the essence of non-identifiability). The bias of the uniform prior to larger inverse dwell times or chemical rates combines in an unfortunate way with the practical non-identifiability problem of the likelihood itself. In contrast, the log-uniform part of the minimally informative prior puts equal statistical weight on each decade and thus alleviates this problem. It would also alleviate the problems mentioned in [35].

Notably, we show that the usually arbitrarily chosen simulation box limits determine the posterior on a relevant scale as soon as the simulation box is large enough given that one uses improper prior distributions. The minimally informative prior desensitizes the posterior concerning the sampling box limits. Only under rare conditions if the posterior has a peak close to the true values, that is multiple orders of magnitude higher than the purely prior-dominated parts, then this problem vanishes. This would make it possible to ignore the strictly prior-dominated parts. However, often, the peak will be less dominant. Importantly, if one uses the minimally informative prior for complex CRNs with a high dimensional parameter space, it is much simpler for the adaptive HMC sampler to produce well-mixing (converging) parameter chains, i.e., the samples indicate that the typical set [93] of the posterior was sampled.

We show that, unfortunately, for typical data qualities and quantities and realistic CRNs, further *objective* or subjective assumptions are necessary to obtain an interpretable and *sufficiently proper* posterior to overcome the challenges from the practical non-identifiability.

A solution to make the posterior proper is to apply meaningful limits to the relevant parameter subset of the sampling box, thereby reducing the uncertainty. The solution is objective if the limits can be theoretically derived (or are rooted in the physical properties of the molecules). Herein, it is shown that this information fusion from data and prior knowledge creates meaningful inferences with the lowest tested data quality, even for the most complex tested CRN. Nevertheless, derived theoretical limits might only sometimes be at hand, or even after their application, practical non-identifiability problems might remain in parameter dimensions where the upper limits do not apply. Herein, an additional vaguely informative prior on the ratio of some rates -a hyperparameter corresponding to the cooperativity of ligand binding and respective unbinding - is applied. Combing these objective and common sense (biochemical) prior assumptions deliver the best inferences in terms of RMSE and uncertainty of the posterior. This additional prior biases the CRN gently towards CRNs where ligand binding and unbinding of the different channel subunits occur independently but still allow for positive and negative cooperativity over orders of magnitude (depending on the choice of hyperparameters). Hence it is a much less radical assumption compared to the commonly used non-cooperativity assumption [87, 88]. Not using such a prior would mean that one is willing to accept *a priori* any order of magnitude of cooperativity effects to occur, which is against commonbiochemical experience - i.e. experience-based priors. Thus, extreme effects are only considered if the data is very certain about them. Thus, this prior is an Occam’s razor.

Using this prior, even without the physically motivated upper sampling box limits, renders the posterior always proper at least in the relevant dimensions of the parameter space since the prior itself is proper. Notably, using these prior assumptions, one can learn from the HMM inference about negative cooperativity within the CRN with at least 10^3^ times smaller data sets than with plain uniform prior assumptions or ML inferences. The allowed reduction of the data quality by a thousandfold is a prerequisite for inferring HMMs of this complexity scale with real world data.

One could also apply this technique to heteromeric proteins or across homomeric proteins containing mutated binding sites [94–96] if the scale of the differences between binding pockets can be coarsely estimated *a priori*. The more coarse the *a priori* estimate is the heavier should the tails of the regularization prior be. A Cauchy prior on the logspace provides a heavier tail but is still a proper prior. A detailed study of the different possibilities is out of the scope of this paper.

In a summary, Bayesian inferences provides flexible tools to accommodate for the shortcomings of ML inferences due to omnipresent practical non-identifiability problems of the likelihood. Careful prior elicitation by being minimally informative where one has absolutely no information on the scale of parameters, but being vaguely informative where there is some physical common sense knowledge about some parameters and very informative when there is *objective* information such as a theoretical upper bound is key get the most of biophysical meaningful HMMs. Using this prior elicitation approach, we where able to obtain meaningful biological insight with a thousand fold lower data quality.

## ACKNOWLEDGMENTS

The authors are grateful to E. Schulz for designing software to simulate channel activity and to Th. Eick for performing simulations. This project was funded by the German Research Foundation (DFG) within Research Group FOR 2518 DynIon (project P2). F. Paul acknowledges funding from the Yen PostDoctoral Fellowship in Interdisciplinary Research and from the National Cancer Institute of the National Institutes of Health (NIH) through Grant CAO93577. M. Habeck acknowledges the Carl Zeiss Foundation funding within the program “CZS Stiftungsprofessure.” We want to thank M. Bücker for helping with the computer cluster at Friedrich Schiller University and Jena and K. Benndorf for their comments on the manuscript.

## 1. Detecting structurally non-identifiable Problem by the smallest eigen value of the Hessian matrix

A heuristic to numerically test for potential structurally non-identifiable likelihoods is to check for the smallest eigenvalue of the Hessian matrix (curvature) of *p*(𝒴_*T*_ | ***θ***_ML_) [97]. If the smallest eigenvalue is effectively zero, then the likelihood is structurally non-identifiable (or almost structurally non-identifiable). Unfortunately, this is only a local criterion for structural identifiable that does not go beyond properties at ***θ***_ML_ and its neighborhood. It indicates based on local information the least confined direction in parameter space. However, the global shape of the likelihood might be more complex [34, 98].

In contrast, strategies such as profile likelihood [33] aim to detect (structurally non-identifiable/practical non-identifiability) pathologies. If *p*(𝒴_*T*_ | ***θ***) is structural identifiable and has only a small degree of practical non-identifiability, the profile likelihood method can construct asymmetric confidence intervals extending the reliability compared to standard ML. Nevertheless, this practice still relies on the assumption of asymptotically large amount and quality of data.

## Appendix A: The practical non-identifiability problem with other states being observed

If a second orthogonal observable is added, such as *y*_2_(*t*) ∝ 𝔼[*n*_*B*_(*t*)], this intrinsic practical non-identifiability problem for *k*_1_ disappears for this CRN but not for *k*_2_ because for the general solution

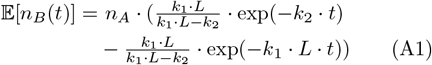

including the mentioned initial conditions assuming *k*_1_ · *L* ≫ *k*_2_ produces 𝔼[*n*_*B*_(*t*)] = *n*_*A*_ · (exp(− *k*_2_ · *t*) − exp(− *k*_1_ · *L* · *t*)). Hense, both exponential decays are preserved. However assuming *k*_2_ ≫ *k*_1_ · *L* for all *L* eliminates both amplitudes, which is an intuative result because **B** is hardly populated. Observing only 𝔼[*n*_*B*_(*t*)] has no practical identifiability problem in the large *k*_1_ direction but still in the large *k*_2_ direction. Having two signals *y*_1_(*t*) ∝ 𝔼[*n*_*O*_(*t*)] and *y*_2_(*t*) ∝ 𝔼[*n*_*A*_(*t*)] resolves the practical non-identifiability for *k*_1_ but does not resolve it for *k*_2_, since

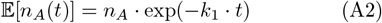

holds. Note, that Eq. A2 has no dependency concerning the rates in the amplitude.

To eliminate the pathologies of the likelihood where *k*_2_ ≫ *k*_1_ holds in ***θ***, one needs to observe data from a mixed population **n**(*t*_0_) = (*n*_*A*_, *n*_*B*_, 0) and a signal that contains two orthogonal observables *y*_1_ ∝ 𝔼[*n*_*A*_(*t*)] and *y*_2_ ∝ 𝔼[*n*_*B*_(*t*)] or alone *y* ∝ 𝔼[*n*_*O*_(*t*)] because 𝔼[*n*_*O*_(*t*)] contains with these initial conditions, in this case, both time scales. Note that because of the initial conditions, the solution of the CME is not a multinomial distribution anymore but a convolution of two multinomials. In other words, the group of ion channels starting in **A** have a different *p*_*O*_(*t*) compared to the group of ion channels starting in **B**.

## Appendix B: Sensitivity analysis of the solution of the RE and the

In chemical physics, it is a well-known phenomenon that often the overall rate of a process in a CRN relies heavily on the slowest chemical reaction, called the rate-limiting step. Rate limitations in inverse modeling become a relevant problem due to the aggregation of Markov states in the signal. Ideally, the mean of an observable of a Markov system would not be aggregated, which means it would report the entire **p**(**K**, *t*) = 𝔼[**n**(*t*)]*/N*_ch_ the population probability vector of all Markov states. In that case, rate limitation would be a minor problem because the kinetics of each state alone allow one to dissect better the contributions of each chemical rate. Only the bandwidth of the data, the signal-to-noise ratio, and the quantity of data would limit the precision of the inference, given a correctly specified likelihood. However, one faces 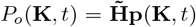 as the observable in PC data, with 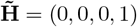 given CRN1 (Fig. 2). Due to the projection, 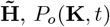 can be *locally* insensitive to changes in some *k*_*ij*_. That means that

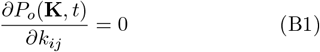

for all *t*.

Instead of *locally*, the *global* 1d sensitivity is scanned (Fig. 10) by scaling in each panel one of the *k*_*ij*_ by [10^1^, 10^2^, 10^3^] and observing the changes of *P*_*o*_(**K**, *t*) for all *t* while leaving the other parameters unchanged. Additionally, some finite experimental time is assumed. Note that multi-dimensionality must be accounted for in a full sensitivity analysis. However, for intuition, it seems sufficient to consider one-dimensional scalings. The deterministic *P*_*o*_(**K**, *t*) of CRN1 (Fig. 2) can be calculated by the RE approach [99]. Qualitatively, the larger the changes of *P*_*o*_(**K**, *t*) compared with *P*_*o*_(**K**_*true*_, *t*) of the CRN (Fig. 10, solid lines), that was used to simulate the data; the smaller can be the data quality to infer that parameter with a satisfactorily small uncertainty, given a correctly specified likelihood. Note that in the following, a likelihood will be constructed [99] for unrealistic best-case data scenarios, whose essential parameter to predict data is *P*_*o*_(**K**, *t*). Thus, differences in *P*_*o*_(**K**, *t*) for each scaling of the *k*_*ij*_, at least for some *t*, are crucial that the model does not have some degree of practical non-identifiability. Even in that best-case scenario, one encounters a practical non-identifiability problem.

**FIG. 10.**
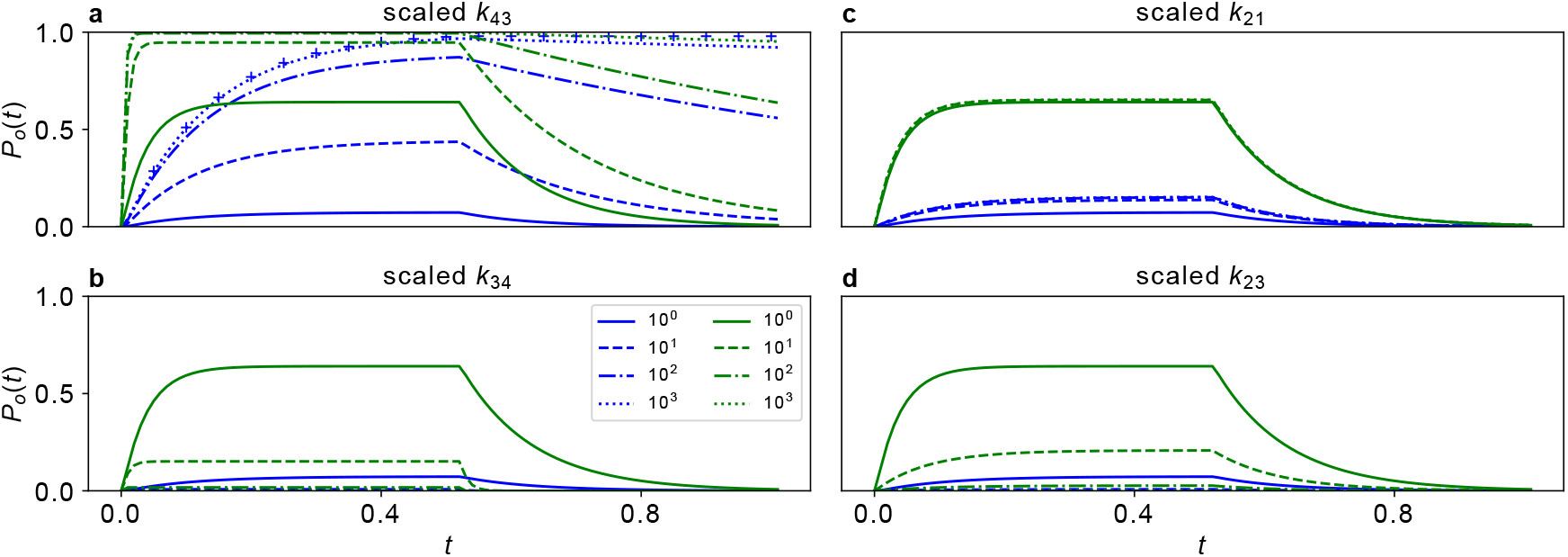
Scaling the chemical rate constants of CRN1 shows that the influence on the mean signal fades, particularly when scaling the ligand-binding rate. In each panel, one chemical rate constant *k*_*ij*_ is scaled by [10^1^, 10^2^, 10^3^] to display the sensitivity of the mean kinetic time evolution of *P*_*o*_(**K**, *t*) to that parameter scaling. Modeling *P*_*o*_(**K**, *t*) by a deterministic RE is identical to experimenting with a noise-free experimental setup that probes an infinite number of ion channels. Two different ligand concentrations [0.125, 1]*µM* are encoded in blue and green, respectively. The different lines indicate the different scaling. In panel **a**, a 10^6^ scaling (blue crosses) for the lower ligand concentration is included. It overlaps with 10^3^ scaling. Indicating that the sensitivity of kinetics ultimately also fades for *k*_43_.

*P*_*o*_(**K**, *t*) is very sensitive (Fig. 10 **a**,**c**) to the two gating rates *k*_43_ and *k*_34_, given CRN1. Firstly, because *P*_*o*_(**K**, *t* = ∞) of both of them can be scaled from 0 to 1 by scaling them from zero to infinity, respectively. Note that for visual clarity (Fig. 10 **a**,**c**) only scaling to larger values is considered. Secondly, both are rate-limiting in the time trace’s respective activation and deactivation parts. However, the experiment’s finite time resolution and duration add global sensitivity limits caused by rate-limiting effects. This can be seen for the smaller ligand concentration *L* = 0.125 *µ*M, the scaling of *k*_43_ (Fig. 10 **a**, blue curves) leads to four visually different kinetics, indicating high sensitivity. However, adding a fourth scaling 10^6^ (blue crosses), the third and the fourth are visually identical, although they are different by three orders of magnitude. For at least the applied 10^3^ scaling 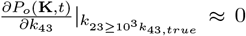. That means the scaled *k*_43_ is instantaneous relative to the other involved transitions. Any higher scaling has little effect on the kinetics of the observable in the given amount of experimental time. However, changes would be detectable in the forward model for an infinite amount of relaxation time because *P*_*o*_(**K**, *t* = ∞) changes with each scaling. Thus, predictions of the model, *P*_*o*_(**K**, *t*), are rate-limited by the other *k*_*ij*_ when *k*_43_ *>* 10^3^ · *k*_43,*true*_. Despite, being locally, around *k*_43,*true*_ *P*_*o*_(**K**, *t*), highly sensitive, the global sensitivity is constrained. From the experimental perspective, that sounds paradoxically because *k*_43_ and *k*_34_ are around *k*_43,*true*_ = 50 and *k*_34,*true*_ = 10 locally the rate-limiting steps themselves for the activation and deactivation time series (unless *L* is very small in the activation data).

These rate-limitations are practically irrelevant for *k*_43_ and *k*_34_ because they are roughly of the order 10^3^ away from their true value, *k*_43,true_ and *k*_34,true_. However, in principle, they exist even for *k*_*ij*_, which are locally around *k*_*ij*,*true*_ the rate-limiting step, or at least for a set of time traces very influential on *P*_*o*_(**K**, *t*).

The global sensitivity of *P*_*o*_(**K**, *t*) changes are *visually* distinguishable in two ways when the scaling is applied on the ligand binding rate *k*_21_ (Fig. 10 **c**). Firstly the sensitivity of *P*_*o*_(**K**, *t*), is reduced (without any effects from the boundary at *P*_*o*_ = 1) for two reasons For *P*_*o*_(**K**, *t* = ∞), the scaling reduces the probability to be in *C*_1_ more and more, but the ratios between the other states *C*_2_, *C*_3_, and *O*_4_ remain the same; hence, the sensitivity of amplitude of the current is constrained from above to smaller values than *P*_*o*_(**K**, *t* = ∞) = 1. Also, *P*_*o*_(**K**, *t*) loses its sensitivity due to rate-limitations; however in this case by the actual transition *k*_43_, which is, from the experimental perspective, the rate-limiting step of the activation. Secondly, for the higher ligand concentration (Fig. 10 **c**, green curves), one can only distinguish 10^0^ and 10^1^ but not the other scalings. For the lower ligand concentration (Fig. 10 **c**, blue curves) one can at least distinguish 10^0^ and 10^1^ and 10^2^ visibly. In contrast to the scaling of *k*_43_ (where the larger *L* has a higher global sensitivity), the smaller *L* reveals a higher global sensitivity of *P*_*o*_(**K**, *t*) for scaling *k*_21_. The smaller *L*, the larger *L* · *k*_21_ can be scaled without being instantaneous relative to the other *k*_*ij*_ in the kinetic scheme. Note that scaling *k*_21_ to smaller values would have no global sensitivity limit as *P*_*o*_(**K**, *t* = ∞) would approach 0. Further at some point *k*_21_ would be the rate-limiting step in the activation for any *L* but this time in contrast to *k*_43_ up the finite limit of the used relaxation time is not even theoretically relevant.

One of the advantages of fitting multiple concentrations (Fig. 10 **a**, green curves) of ligands (or, depending on the molecule, other series of stimuli) that the limiting values for *k*_*ij*_, above which rate-limitations eliminate all effects of altering *k*_*ij*_ on the likelihood can be tweaked by changing the magnitude of the stimulus. Hence, the local and global sensitivity of *P*_*o*_(**K**, *t*) to altering some *k*_*ij*_ is changed.

Similarly, *P*_*o*_(**K**, *t*) has a high sensitivity for scaling up the ligand-unbinding rate *k*_23_ (Fig. 10 **d**) because the scaling pushes *P*_*o*_(**K**, *t*) → 0 for all *t*.

Summing up, there is a global sensitivity limit of the mean kinetics and equilibrium to scaling up *k*_21_ and similar *k*_32_. However, for scaling *k*_43_ up, the global sensitivity limit also requires a finite relaxation time in the experiment. However, a global sensitivity limit for scaling up *k*_34_, *k*_23_, and *k*_12_ does not exist. And similarly, scaling one of the down *k*_21_, *k*_32_ and *k*_43_ will decelerate the coupled reaction more and more and *P*_*o*_(**K**, *t* = ∞) will approach 0 such that the sensitivity of *P*_*o*_(**k**, *t*) ≈ 0 to scaling these rate constants to smaller values is high. These asymmetries in the sensitivity will produce highly asymmetric likelihood profiles (Fig. 11). If one had one independent observable for each Markov state, meaning that the full mean **p**(**K**, *t*) is an observable (i.e., no aggregation of states in the fictitious signal); the limitations due to partial observability would be alleviated. In this optimal case, only the signal-to-noise ratio and the bandwidth would induce practical non-identifiability structure in the likelihood. Adding orthogonal signals fluorescent signals such as done with cPCF [100] is the attempt to expose more of **p**(**K**, *t*) than *P*_*o*_(**K**, *t*) does.

**FIG. 11.**
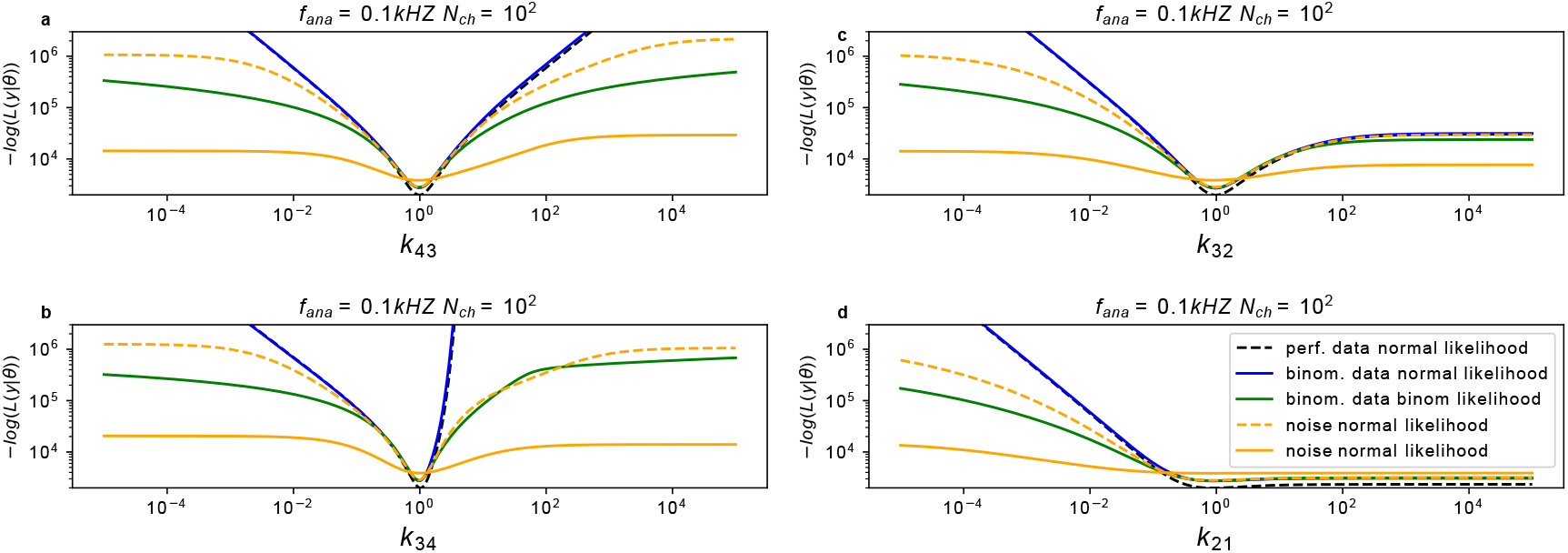
Likelihood Profiles of four different parametersof CRN1 reveal varying degrees of practical non-identifiability even with instrumental noise-free data. Here, we show likelihood profiles, given different data settings and corresponding likelihoods for different *k*_*ij*_. **a-d**, The black dashed curve indicates the value of a normal log-likelihood analyzing the perfect data (no experimental and intrinsic noise). The perfect data are the result of calculating the deterministic solution of the RE. The blue line represents binomial data analyzed with a normal likelihood. Green represents the binomial data analyzed with binomial likelihood. We add white normal distributed noise to the binomial data (orange curves) with *σ* = 1 (dashed curve) *σ* = 10 (solid curve).

### 1. HMMs, even under unrealistic optimal data and prior knowledge scenarios, have a degree of a practical non-identifiability problem

The following section scrutinizes the likelihood of PC data of the 4-states-1-open-state model for unrealistic optimistic data and prior knowledge scenarios. It is demonstrated that in these best-case scenarios, the likelihood has a degree of a practical non-identifiability problem, which indicates that HMM inference has, in general, a practical non-identifiability problem.

A practical non-identifiability problem exists due to an infinite area in ***θ*** where the likelihood is insensitive to changes of some *k*_*ij*_’s or 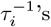. The reasons for the non-sensitivity are further investigated in App. B by comparing the deterministic mean kinetics, *P*_*o*_(**K**, *t*), for different **K**. The combination of rate-limitations with the limited influence of some *k*_*ij*_ on the equilibrium open probability, *P*_*o*_(**K**, *t* = ∞), causes *P*_*o*_(**K**, *t*) to lose its sensitivity towards changes of the chemical rates in an infinite volume in ***θ***. One might consider it as intrinsic inferential limits of the protein’s model likelihood. Inferring some of the *k*_*ij*_’s with a low uncertainty defines for the other more uncertain *k*_*ij*_’s where the likelihood becomes completely insensitive in ***θ*** for the value of these *k*_*ij*_’s. where in ***θ*** the likelihood becomes completely insensitive for these *k*_*ij*_’s. Ultimately, this causes the likelihood of partially observed CRNs to have a degree of practical non-identifiability even for completely experimental noise-free data.

### 2. The fading sensitivity of some chemical rates on the observed kinetics induces practical non-identifiability

When modeling experimental data, the intrinsic inferential limits interact with the data’s bandwidth, experiment duration, and signal-to-noise ratio. Nevertheless, it is shown herein that the intrinsic inferential limits of a CRN might become the bottleneck for the degree of practical non-identifiability for many *k*_*ij*_’s or 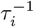 s the entire model.

In the ML context, it is one prerequisite for entire practical identifiability in contrast to practical non-identifiability [35] that the parameter values of likelihood surfaces of constant value form compact sets. In a 1D setting, one could formalize this prerequisite to that

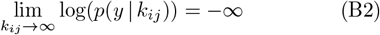

holds. Similarly,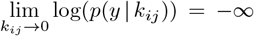. In other words, the model’s predicted statistics, given the set of parameters, have to be further and further away from the true statistics of the data if some parameters are more and more diverging. When sampling a posterior, given a minimal informative prior, with a finite sampling box, one assumes that Eq. B2 is sufficiently fulfilled at the limits of the sampling box.

The discussion of the forward model (App. B) indicates that the likelihood of macroscopic HMM data beyond trivial examples has the opposite of Eq. B2 as a general feature: If the global sensitivity of *P*_*o*_(**K**, *t*) for some scaling magnitude is vanishing, then the derivative of the log-likelihood must approach 0.

In the following, a simulated best-case data and prior knowledge scenario shows that practical non-identifiability features arise in a likelihood which is an exact representation of the statistics of the simulated data. The practical non-identifiability problem becomes essential for maximum likelihood estimation or Bayesian statistics at poor data quality. Nevertheless, it is always present in principle. Again, this does not mean there is no optimal parameter set in the posterior or likelihood, but the corresponding uncertainty quantification faces severe problems (Fig. 1). Suppose the likelihood of the model used for the inference has a unique global optimum but becomes flat in some directions. In that case, the inference quality must be judged by how high the likelihood’s maximum is relative to its flat area. In the case of Bayesian statistics with the applied Jeffreys prior, one can transform this criterion into how much the posterior decays (5 **a**,**b**) before its slope is approximately the slope of the log-uniform prior. Corresponding to the profile likelihood technique [33], one may define that, for example, 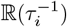 needs to be decayed to 10^*−*2^ relative to the value of the posterior’s peak before its slope drops to 1.05 of the log uniform prior slope to call the parameter identified. Then *N*_ch_ = 10^2^ (5 **a**) would be by definition not be considered identified, even if the posterior has some interval which is clearly distinguished from the flat rest of ***θ***. In contrast, if we define that the posterior needs to decay at least to 10^*−*1^ before its slope drops to 1.05, then it would be considered as identified. One needs this definition based on how high is the peak of the posterior relative to the prior dominated area to justify the limits of the sampling box after one has done the sampling. That is because outside of the sampling box is always an *infinite amount of probability mass* due to the practical non-identifiability problem, given that an improper prior (Eq. 8) is used.

### 3. The profile likelihood reveals flat parts even in a best-case scenario

We generate idealized model data sets from CRN1 (Fig. 2) and then add in a step-wise manner intrinsic fluctuations of the ensemble and instrumental noise to the synthetic signal. We calculate for each time point the open probability *P*_*o*_(**K**, *t*) of the model and derive from it the mean signal

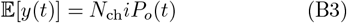

and its variance

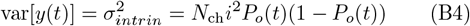

utilizing the multinomial approximation of the distribution of stochastic gating and binding of ion channels in macroscopic patches [99]. The initial preparation of the relaxation experiment assumes that all channels are at *C*_1_ because the ligand concentration holds *L* = 0. Further, only one data point per time trace is used per activation and relaxation experiment With these conditions, the multinomial assumption used in [99] precisely describes the data’s statistics. Otherwise, the actual likelihood, given the Markov assumption, must be based upon a convolution of multinomial distributions [38, 77].

As asymptotically ideal data, we use the true model’s mean current signal (Eq. B3) without any intrinsic or extrinsic noise (Fig. 11, black dashed curves). Then, we additionally sample the stochastic time evolution by drawing from a binomial distribution *n*(*t*) ∼ binomial(*N*_ch_, *P*_*o*_(**k**, *t*)) current data for each time step (Fig. 11, blue and green curves) and finally add some experimental white noise with constant variance (Fig. 11, orange curves). In that way, we generate data sets that match the white noise assumption of the RE approach [99], which differs from simulating data by the Gillespie algorithm. In contrast to the Markov property (i.e., included in the Gillespie algorithm) of actual experimental time series data, we switch the MM property off. The MM property lowers the information content of a time trace (see, [99] Fig. 10). Hence, the used data sets contain more information than actual HMM data with the same sampling frequency. Consequently, the following analysis is a best-case data scenario corresponding to a massive bootstrapping of the data: one data point per time series (see, [99] Fig. 10).

These data sets are analyzed by assuming different likelihoods. For the perfect data set (Fig. 11, black dashed curves), we use a normal log-likelihood which inherits its mean signal and variance from the above-stated multinomial assumption of the RE approach [99]. The blue curve represents the log-likelihood value for binomial data analyzed with a normal log-likelihood and, correspondingly, analyzed with a binomial log-likelihood (green curve). We fix all parameters but one to their true values and plot the profile log-likelihood vs. the scaled 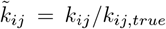 parameter. This way, a very informed prior knowledge of the rate matrix is assumed, as every other relevant parameter except the scaled one is precisely known. Our inverse modeling scenario corresponds to the forward modeling scenario discussed in (Fig. 10). In contrast to [33, 36, 101], the other parameters are not optimized such that the profile likelihood herein displays the contour along a straight line in ***θ*** while their procedure searches for the slowest decay of the likelihood given one fixed parameter as a constraint, which is not necessarily a straight line; it is some lower dimensional, potentially curved submanifold in ***θ***. However, the approach used herein seems sufficient for displaying pathologies of the 1D likelihood. Note that the profile of the log-likelihood is identical to the profile of a 1D log posterior derived from a uniform prior distribution. Nevertheless, if more chemical rates are unknown, likelihood profiles are generally not identical to the 1D marginalized posterior.

In the first column (Fig. 11 **a**,**b**) we investigate *k*_43_ and *k*_34_ the opening and closing rates of the channel, respectively. Even if it is not displayed, there exist inevitably an upper limit of − log(*p*(*y* | *k*_*ij*_)) for scaling *k*_43_ because of the observed loss of sensitivity of *P*_*o*_(**K**, *t*) “occurring” above scalings of 10^3^ (Fig. 10 **a**). That is because the normal likelihood uses only the mean signal and its variance as parameters, which are functions of *P*_*o*_(**K**, *t*). Ultimately, *P*_*o*_(**K**, *t*) becomes insensitive even to changes of large *k*_43_ and *k*_34_ (Fig. 10 **b**, green curves). Hence − log(*p*(*y* | *k*_*ij*_)) cannot go to infinity. Correspondingly to the large interval, in which *P*_*o*_(**K**, *t*) is sensitive (Fig. 10 **a**) the limiting behavior of − log(*p*(*y* | *k*_*ij*_)) is beyond the limits of panel Fig. 11 **a**.

Note that the more accurate binomial log-likelihood reveals a slower increase of its negative log-likelihood values (Fig. 11 **a-d**). This means that the normal approximation rules out large deviations from the true value more strictly than the true likelihood. The difference between the binomial and normal log-likelihood is reminiscent of the breakdown of the normal approximation to the binomial distribution when *P*_*o*_(**K**, *t*) is close to 1 or 0. The vanishing sensitivity of *P*_*o*_(**K**, *t*) of other rate constants for *k*_34_ and *k*_43_ are (Fig. 11 **a**,**b**), in the considered case, not the bottleneck as additional experimental noise (Fig. 11 **a** orange curves) strongly affects the depth of the minimum. If we add some noise with *σ*_*exp*_, the total variance of the signal will be 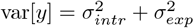. With a small amount of extra experimental noise (Fig. 11) **a**,**b**, orange dashed curves), the likelihood profile for *k*_34_ and *k*_43_ becomes flat, giving each of the parameters a more noticeable degree of practical non-identifiability. As expected, a larger magnitude of the noise (orange, solid curves) makes the minimum more shallow.

Of course, the resort in practice is that the minimum negative log-likelihood is deep enough. It is still almost an order of magnitude deep, and the minimum is of the order of 10^4^ such that in the space of exp(log(*p*(*y* | *k*_*ij*_))) where one evaluates parameter non-identifiability, the corresponding relative height of the likelihood’s maximum is even higher above the flat parts. That enables almost arbitrary high-confidence thresholds [33] in the probability space. The word ‘almost’ seems not even worth a technical note, given that the data used to calculate the log-likelihood profile creates such a low minimum in panels a and b. Still, it is impossible to choose an arbitrarily high confidence threshold (Fig. 1, black dashed lines) in the maximum likelihood framework that delivers a finite confidence region for the parameter *k*_43_. Bayesian analysis with a Jeffreys prior would technically have the same practical non-identifiability feature, leading to an improper posterior on *k*_43_ ∈ (0, ∞). The practical non-identifiability aspect, induced by the rate limitations due to other finite chemical rates, of the HMM becomes relevant when considering *k*_32_ and 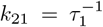 (Fig. 11 **c**,**d**), respectively under identical experimental conditions. The right-hand side of the likelihood is flat even under the perfect data assumption because *P*_*o*_(**K**, *t*) reached its global sensitivity limit (Fig. 10, **b**). In other words, it is more suited for inverse modeling if there is a global sensitivity limit, the likelihood based on a rate matrix with an infinitely scaled *k*_32_ or 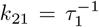 does not become an infinitely bad predictor of the data, but that is what the Eq. B2 requires in order that the model has no degree of practical non-identifiability. Still, if the data quality is enough, the minimum of the negative log-likelihood might be deep enough.

If one extrapolates these observations, one can conclude the following: On the one hand, the more unknown parameters the rate matrix has, the more flexibility might the likelihood have to repeal the effect of one misplaced parameter in the signal by tweaking other parameters; hence, more or higher-dimensional submanifolds might emerge in ***θ*** on which the likelihood profile (surface) is flat. If one needs to learn the full rate matrix, the *k*_*mn*_, which influences the data the most, given the tested conditions, will have the sharpest marginal posterior. That defines where the marginal posteriors of the other *k*_*ij*_ become flat. Because at some value of *k*_*ij*_, the likelihood will have lost all sensitivity to the exact value of *k*_*ij*_. This does not mean that these *k*_*ij*_ do not have a peak somewhere else. On the other hand, it is straightforward to construct with more unknown chemical rates a direction in the higher dimensional ***θ*** where the rate limitation of a single chemical constant by other chemical rate constants is repealed by scaling others, too. Assuming a scaling acting simultaneously on *k*_21_, *k*_32_, and *k*_43_ and that the other chemical rates are fixed at their true value, The scaling means moving the chemical rates along a diagonal in ***θ***. Moving along this direction the likelihood has the desired 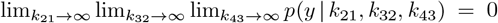 for data, which is uncorrupted from experimental noise. In this limit, the kinetics become a step function from *P*_*o*_(*t* = 0) = 0 to *P*_*o*_(*t* = Δ*t*) = 1. Given the finite bandwidth of real experimental data, this happens already for finite *k*_21_,*k*_32_, and *k*_43_. In contrast, for each direction along the parameter axis *k*_12_, *k*_23_, or *k*_34_ alone 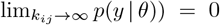. So, we can deduce from the model structure by the forward model at least four trivial directions in ***θ*** for which the likelihood behaves rather PI-like, given ideal data without any experimental noise. The existence of some PI-like directions in the parameter space is, of course, not sufficient to make the model entirely PI.

In conclusion, in a Bayesian framework, the posterior sampled in a sufficiently large sample box with enough samples would always be dominated by the prior in the right tails of the inverse dwell times. Thus, if we try to be uninformative by equipping the time scales with a log-uniform prior and setting the sampling box limits to huge values, we end up with a posterior, which depends accidentally on the nonphysical limits of the sampling box. Strictly speaking, this is true for all parameters. The depth of the log-likelihood minimum around the actual value ensures that this is unnoticed in most data scenarios. That said, posteriors derived from the minimal informative prior still have a peak (Fig. 5**a-b** around the true value and cover the true value within a considerable part of their peak; thus, we still gained information.

Notably, in the strong data, case [38], without knowing that the likelihood becomes flat in the inverse time-like parameters and without any elaborated prior selection, we showed in [38] that the HDCVs have the expected coverage frequency over the true values. In other words, the calculated Bayesian uncertainty of the parameters matches the frequency of finding the parameters within a volume to a high degree if the algorithm correctly maps the first-order Markov property of the data. Thus, cutting away infinite “probability mass” at some data strength is just fine because the rest behaves as if the posterior is well-defined. In contrast, in the weak data case where the posterior drops only a limited ratio before it is “flat,” the limits of the samples are more critical. One could define thresholds that sort out samples in the flat likelihood area and define the posterior from the remaining, equivalent to sampling it with a smaller sampling box. It is beyond the scope of this article to investigate the necessary numerical criterion, what is flat enough that one should consider the part of the posterior as dominated from the prior. The performance of such an approach is also out of the scope of this article.

## Appendix C: Definition of minimally informative (MI) priors by Jeffreys’s rule

### 1. Nonlinear coordinate transformations alter the functional form of a prior

To demonstrate potential problems of uniform priors in combination with weakly informative data and to contrast them against the benefits of a minimally informative prior, a definition of how to construct a minimally informative prior, *p*(**K**), is required. We want the posterior to be as sensitive to the data as possible when little is known about some parameters. At first glance, a uniform prior seems to be the least biased choice. However, uniformity of a distribution is not invariant under non-linear parameter transformations. Therefore, a uniform prior is not in general MI, it often induces biases [63]. For example, one can parameterize a model of a monomolecular chemical reaction in terms of either the rate constant *k* or the activation energy *E*_*A*_. Their relationship is often approximated by Arrhenius’s law 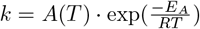, where *R* is the gas constant, *T* the temperature and *A* some temperature-dependent proportionality constant. Let us define *k*_max,box_ as the upper limit of the sampling box. What shape has the prior 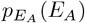 in the parameterization of *E*_*A*_, if we assume *p*_*k*_(*k*) = uni(0, *k*_max,box_) in the *k* parameterization and *A*(*T*) and *T* to be known? It turns out that *E*_*A*_ would be exponentially distributed. However, if one assumes a log-uniform distribution 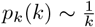, then *E*_*A*_ would be uniformly distributed.

### 2. Different parameters need to have different minimally informative priors

Parameters of many probability distributions can be classified into location, scale and shape parameters based on their behavior under transformation groups. The probability density *p*_*s*_ of a scaling parameter *s* obeys *p*_*s*_(*y*) = *p*(*y/s*)*/s*. Typical examples of scaling parameters are the standard deviation *σ* of a normal distribution or the mean dwell time *τ* to remain in a specific Markov state. Note that dwell times *t*_dwell_ ∼ 1*/τ* exp(− *t*_dwell_*/τ*) are exponentially distributed. Various notions of “minimal information” have been proposed [67], which can result in different priors. For example, Jaynes [67] and Jeffreys [64] argued for a log-uniform prior for a scaling parameter:

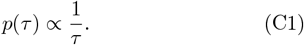

Jeffreys’s arguments are mainly based on a specific non-linear coordinate transformation. He argues that the log-uniform prior is invariant under power transformations, meaning that any *u* = *τ*^*ξ*^ with arbitrary exponent *ξ ≠* 0 has the same functional form

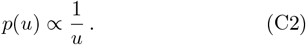

For example, a minimally informative statement about *σ* should be also minimally informative for the variance *σ*^2^. Applied to the parameters of CRNs, the power transformation *k* = *τ*^*−*1^ also identifies a chemical rate constant as a rate parameter. Note that here the term “rate parameter” should be understood in a statistical sense which defines a “rate parameter” as the inverse of a scaling parameter. Thus, the functional form of a minimally informative prior for *k* remains a log-uniform distribution. Jaynes’s arguments [67] are based on transformation groups and lead to Eq. C1, whereas Jeffreys’s original rule yields a different result. In his original formulation [63], Jeffreys defines the prior by

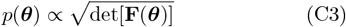

where 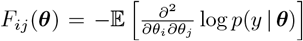 is the Fisher information matrix. Jeffreys’s prior creates a posterior that transforms covariantly under reparameterizations of the model, meaning that the same probability will be assigned to identical volumes in parameter space independent of the parametrization. Despite its success in single-parameter problems, Jeffreys’s rule (Eq. C3) is challenged by multiple inconveniences in multi-parameter inferences [102, 103]. For example, Jeffreys’s rule (Eq. C3) applied to a Normal distribution model with unknown mean *µ* and variance *σ*^2^ results in a prior *p*(*µ, σ*) ∝ *σ*^*−*2^, contradicting Eq. C1 of [67] but also the revised version of Eq. C3 [64]. If, on the other hand, each parameter is treated independently, 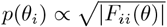, assuming that the others are known, then we obtain the log-uniform prior *p*(*µ, σ*) ∝ *σ*^*−*1^. Thus, we will use a revised version Eq. C3 in the following, which treats location, scale and shape parameters independently, as suggested by Jeffreys himself [64] and consistent with [67]:

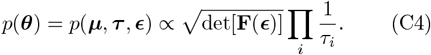

The location parameters, ***µ***, such as the mean value of the normal distribution, are assigned uniform priors. Each scaling parameter, ***τ***, has a log-uniform prior. Only the shape parameters ***ϵ*** are treated conjointly according to Jeffreys’s rule [64]. In addition, we simplify Eq.C4 by assuming that 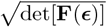 can be applied to each column of **K** (Eq. 16) independently. In that way, we obtain closed-form solutions of 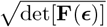 derived from simpler statistical models for the remaining ***ϵ*** of each column of **K**.

## Appendix D: Illustration of the prior for a column of the rate matrix with two Markov transitions leaving the state

Here, we discus the shape of the minimally informative prior corresponding to one column of **K** for two different scenarios. If the CRN assumes, e.g., two existing Markov transitions, *W*_*i*_ = 2, leaving the *i*-th state, the corresponding prior needs to be a beta distribution (2d Dirichlet distribution, red, inset of Fig. 12**a**). The probability drawn from the beta distribution is *ϵ*_*ji*_, and because there is only one alternative transition, the corresponding probability of that transition equates to 1 − *ϵ*_*ji*_. Three assumed Markov transitions, *W*_*i*_ = 3, require a 3d Dirichlet distribution (Fig. 12**a**); four connections are distributed by a 4d Dirichlet distribution, et cetera. The only exception is a Markov state at some CRN’s end. This Markov state is only connected by one transition to the network. In that case, 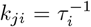 such that there is only the log-uniform distribution (Fig. 12**b**) present for this column of **K**. To understand the difference in the assumptions between a log-uniform distribution for 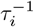 (Fig. 12**b**, blue curve) and a uniform prior on *τ*_*i*_ (green curve) consider the following. A uniform prior states that one has ten times more probability in [10^1^ − 10^2^] than in [10^0^ − 10^1^] that the actual value is in that decade. This relationship continues for each decade. In contrast, if one assumes a log-uniform prior such that the exponent is uniformly distributed, it means that each decade has the same probability. Notably, both distributions, uniform and log-uniform, are improper (Eq. 8), which causes the posterior to be improper if the likelihood is too flat. Similar to the uniform distribution, the log-uniform distribution is only a distribution if restricted to some interval [*a, b*] ⊂ (0, ∞) with 0 *> a > b <* ∞. That has potential risks for any statistical inference. Below, App. B 1, indicates that it is a general feature of HMMs that the likelihood is flat in a space of infinite volume and only in a finite volume around **K**_true_ is the likelihood, not flat. That means using improper priors should lead to an improper posterior, which we can reproduce (Fig. 5). So, careful checking of the posterior’s tails is needed to understand how sensitive the results of the inference are to the typically subjective limits of the sampling box.

**FIG. 12.**
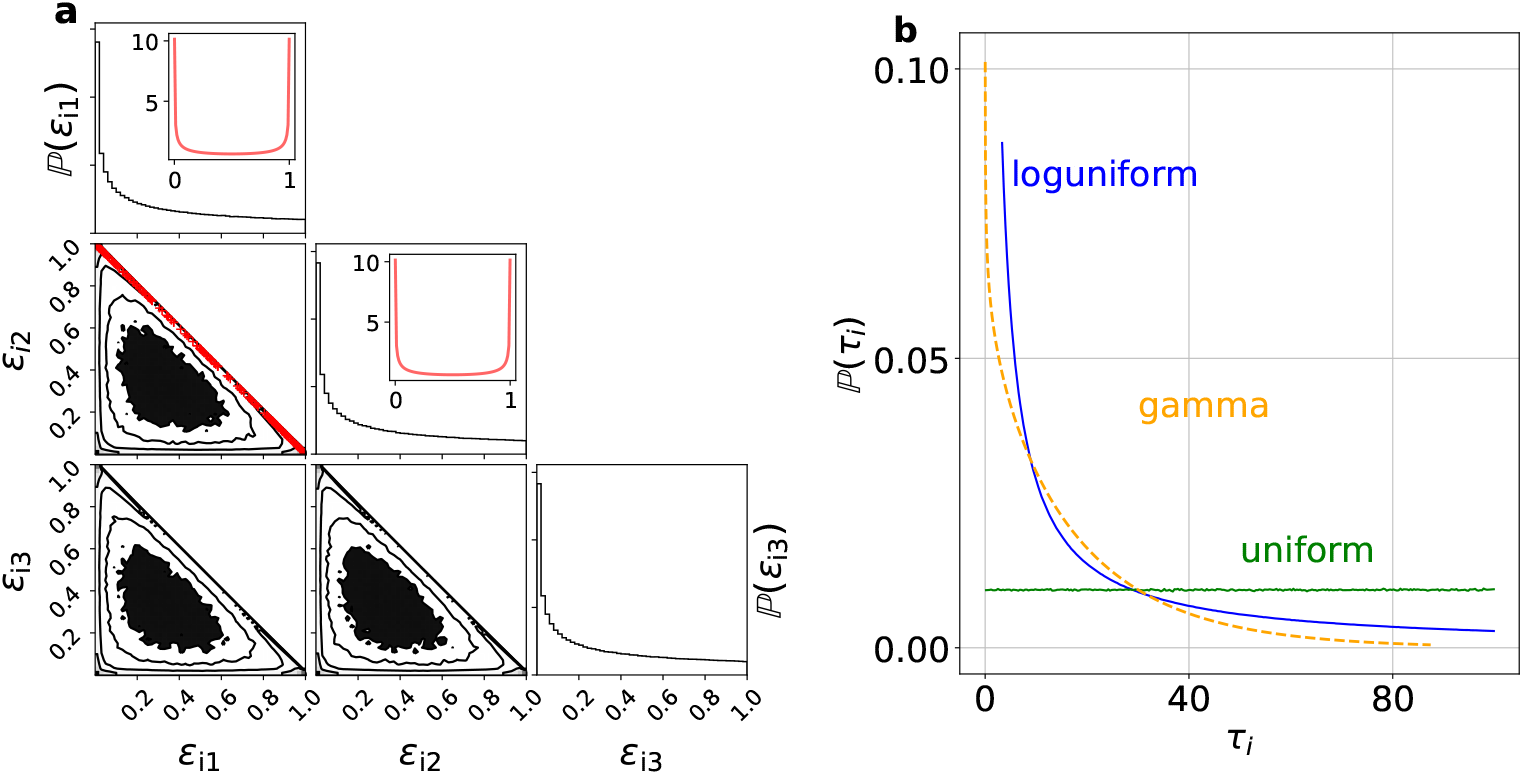
MI prior for a Markov state with supposed three transitions that exit the state. **a** A 3d Dirichlet prior dir(0.5, 0.5, 0.5) (black) for a supposed state which has three Markov transitions leaving the state such as state *C*_4_ in CRN2 (Fig. 2). For a supposed state with two exiting Markov transitions, the corresponding dir(0.5, 0.5) = beta(0.5, 0.5) (red) is displayed in the first two columns. Usually, the beta distribution is only considered as the distribution of success probabilities. Nevertheless, each draw from a beta distribution is a draw from a probability simplex by *P*_success_ and *P*_fail_ = 1 *− P*_success_. The constraints Σ_*i*_ *ϵ*_*i*_ = 1 and 0 ≤ *ϵ*_*i*_ ≤ 1 of the probability simplex in 2d create a 1d submanifold, simply a straight line (red). However, the constraints define a 2D submanifold embedded in 3D when three outcomes exist. **b**, shows the possible alternatives for the prior on *τ*_*i*_ or one *k*_*ji*_ of column the *i*-th state which sets the time scales of the Markov transitions leaving the state. The panel contrasts the log-uniform prior (blue) against the uniform prior (green) and the a gamma prior (yellow) which would have regularising effect if small but finite parameters are used. A simple way to draw from a log-uniform distribution is *τ*^*′*^∼ uniform(*a, b*) followed by 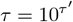. This way of drawing log-uniform random numbers shows that each decade is equally probable.

## Appendix E: Some considerations when the RMSE should be evaluated on the logarithm of the parameters

To calculate the RMSE of *p*(**K** | 𝒴 _*T*_), the samples are reparametrized back to *k*_*ij*_’s for the following reason. The probability, *ϵ*_*ji*_, which Markov transition is chosen, exists on a probability simplex.Relative errors from **K**_true_ cannot be unambiguously merged into the RMSE as long as the boundaries are close. Nevertheless, similar problems arise with chemical rate constants at *k*_*ij*_ = 0. A 10 times larger parameter value contributes with a relative error of 9 according the definition in [38] while a 10 times smaller parameter value contributes only with 0.9 relative error. However, it is reasonable to care more for the multiplicative deviation 𝔼[*k*_*ij*_*/k*_*ij*,true_] because *k*_*ij*_’s are, in the statistical sense, inverse scaling parameters. When the information in the data is weak, one expects large multiplicative deviations from **K**_true_ and correspondingly a diffuse *p*(**K** | 𝒴 _*T*_), which are influenced by the mathematical constraints. *p*(**K** | 𝒴 _*T*_), will be far away from multinomial distributions. Hence, a redefinition (Eq. 21) compared to the definition of the RMSE of [38], seems obligatory. The boundaries at zero are eliminated by log transformation. The mean value is the standard point estimator in Bayesian statistics. It converges towards the peak of *p*(**K** | 𝒴 _*T*_) if the distribution becomes monomodal and symmetrical. Nevertheless, it is shown (inset, Fig. 5) that reporting the peak of the marginal distributions due to the practical non-identifiability problem makes sense. Note that for small multiplicative deviations (i.e., 𝔼[*k*_*l*_]*/k*_*l*,true_ ≈ 1) and using a Taylor series expansion, one can show that

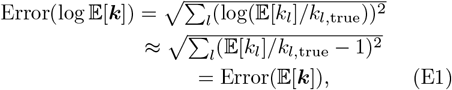

holds. Hence, the redefinition becomes equivalent to the of the RMSE in [38]. In aid of illustration, we use the PC data (Fig. 2) and compare Error(𝔼[***k***]) of the definition in [38] with the new definition. As previously done, *N*_ch_ is used as a measure for the PC data quality (Fig. 13); thus, also as a proxy for the information contained in the data for a given model. The transition (Fig. 13) that both definitions become equivalent occurs roughly at *N*_ch_ *<* 2 · 10^3^ = *N*_ch,crit_. Below *N*_ch,crit_, the RMSE of the logspace is compared to the error on chemical rate space 1 − 2 orders of magnitude smaller. Only minor differences can be distinguished above *N*_ch,crit_.

**FIG. 13.**
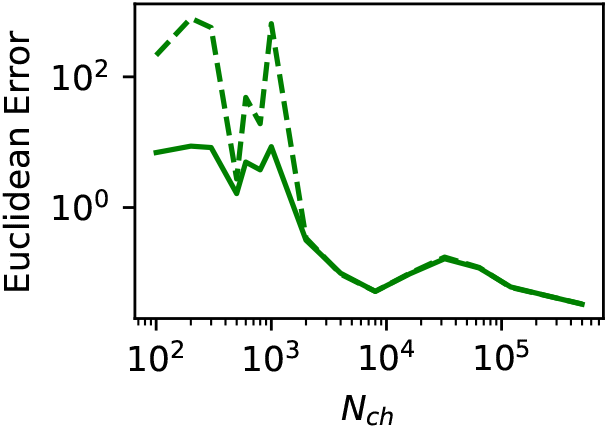
Comparing the two RMSE definitions based on the previously used CRN1 PC data. The solid line corresponds to the RMSE on the log space. The posterior was sampled using a uniform prior. The two RMSE definitions are equivalent for *N*_ch_ ≥ 2 · 10^3^. The two definitions deliver indistinguishable results at *N*_ch,crit_.

### a. The learning of the model on the non-log parameter space

The influence of the Jeffreys rule inspired prior with diffusion limit on the single parameter level (Fig. 14) is visualized by HDCIs (Eq. 12) vs. *N*_ch_. For clarity, we plot for the uniform prior (green dashed lines) and for the Jeffreys prior (gray area) only the 0.95-HDCI. However, a series of HDCIs is plotted for the diffusion limit prior (blue area). The log-space is not used; instead, a log-log scaling is applied to indicate the skewness of the marginal posteriors. Due to the skewness of the marginal posterior distributions, the HDCIs differ from quantiles. The posterior of 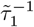, which is also identical to 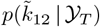 (blue posterior), since 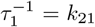 is cut by the diffusion limit unless the data quality is very high. The additional information allows confining the samples to a much more restricted space producing decent inferences even at *N*_ch_ = 10^2^. The bias towards too large 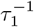 of 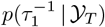 using the minimally informative prior now turns into a bias to too small 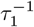. The applied constraints on *k*_32_ have also for 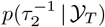, to which the second binding rate *k*_32_ contributes by *τ*_2_ = *k*_32_ + *k*_12_, a regularizing influence.

**FIG. 14.**
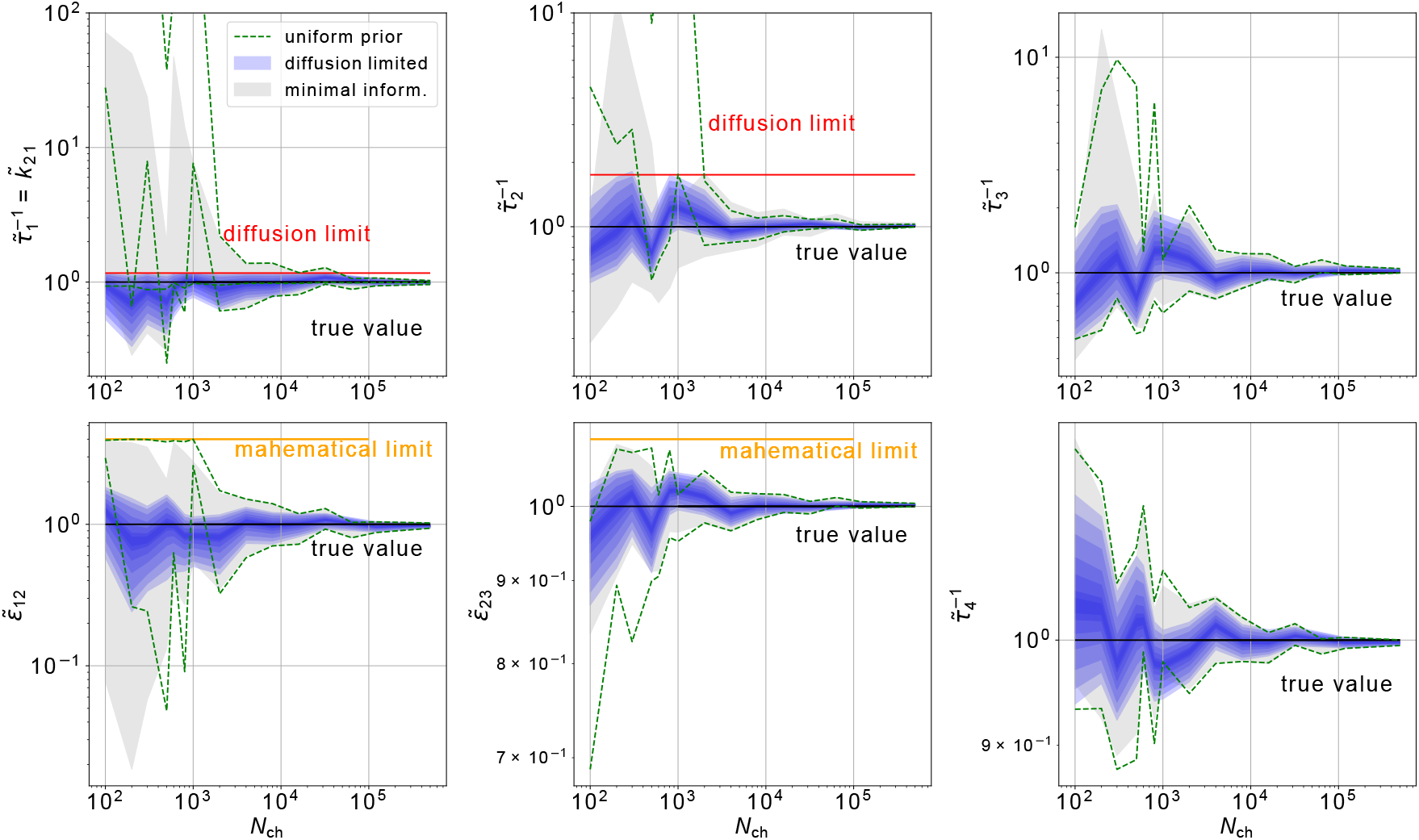
The marginal posterior quantiles of CRN1 vs. *N*_ch_ **displays an influential diffusion limit even for high** *N*_ch_. The red horizontal bars indicate the physical diffusion limit applied in sampling the blue posterior. The black horizontal bars indicate the true values. The orange horizontal bars indicate the mathematical limit that probabilities hold *<* 1. Due to the scaling to each true parameter, this limit is ≠ 1. The green marginal is a 0.95 HDCI with a uniform prior over **K**. The gray area indicates the 0.95-HDCIs using the minimally informative prior with no diffusion limit applied. The set of posterior [0.95, 0.9, 0.8, 0.6, 0.4, 0.2, 0.1]-HDCIs using the minimally informative prior with enforced diffusion limit is shown in blue. Due to the structure of the CRN (only one transition leaves *C*_1_), 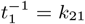 holds. Hence, the diffusion limit is identical on both parameters. For thesecond ligand binding *k*_32_, *k*_32_ ∼ log-uniform(k_32_) is modeled, and the diffusion limit is enforced on this parameter.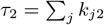 is shown, a probability mass above the red bar is possible.

That is true even though it cuts only through the far tail *p*(*k*_32_ | 𝒴 _*T*_) (not shown here). In the regime *N*_ch_ ∈ [10^2^, 10^3^], the constraint makes a very vague *p*(*τ*_1_ | 𝒴 _*T*_) where the 0.95-HDCI covers up to two orders of magnitude, including a potential practical non-identifiability problem, to a decent *p*(*τ*_1_ | 𝒴 _*T*_). The 0.95-HDCI spans [0.4, 1.16] in units of 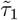. The highly skewed distributions justify the redefinition of the Euclidean error, the log space.

## Appendix F: More data sets for CRN2 showing the gain of robustness by the minimally informative prior

We continue investigating the benefits and challenges of the minimally informative prior (continuing the discussion of Fig. 7) arising when the complexity of the CRN increases. In Fig. 15 **a** the pathological dimensions of *p*(**K** | 𝒴 _*T*_) derived from the uniform *p*(**K**) (colored) against the same dimensions of *p*(**K** | 𝒴 _*T*_) (black) derived from the minimally informative *p*(**K**) are compared. The less pathological dimensions in the sense that they deliver roughly Gaussian marginal posteriors are plotted in Fig. 15 **b1-6**. We reduce the data quality for the minimally informative prior (to create a more challenging scenario), showing the overall gain in the robustness of the inference by the minimally informative prior. We further have to work with smaller sampling boxes (which still contains **K**_true_) for the uniform prior (because of an often not converging sampler). Using a smaller sampling box for the uniform prior also disadvantages the minimally informative prior, see Arguments in Sec V F. Thus, the *sufficiently proper* posterior for *N*_ch_ = 5 · 10^5^ using the minimally informative prior, shows the overall robustness gain due to the minimally informative prior. For *p*(**K** | 𝒴 _*T*_) based on a uniform *p*(**K**) we use a set of data sets with practically *impossible* to achieve data quality *N*_ch_ = [7.5 · 10^4^, 10^5^, 2.5 · 10^5^] for PC experiments.

**FIG. 15.**
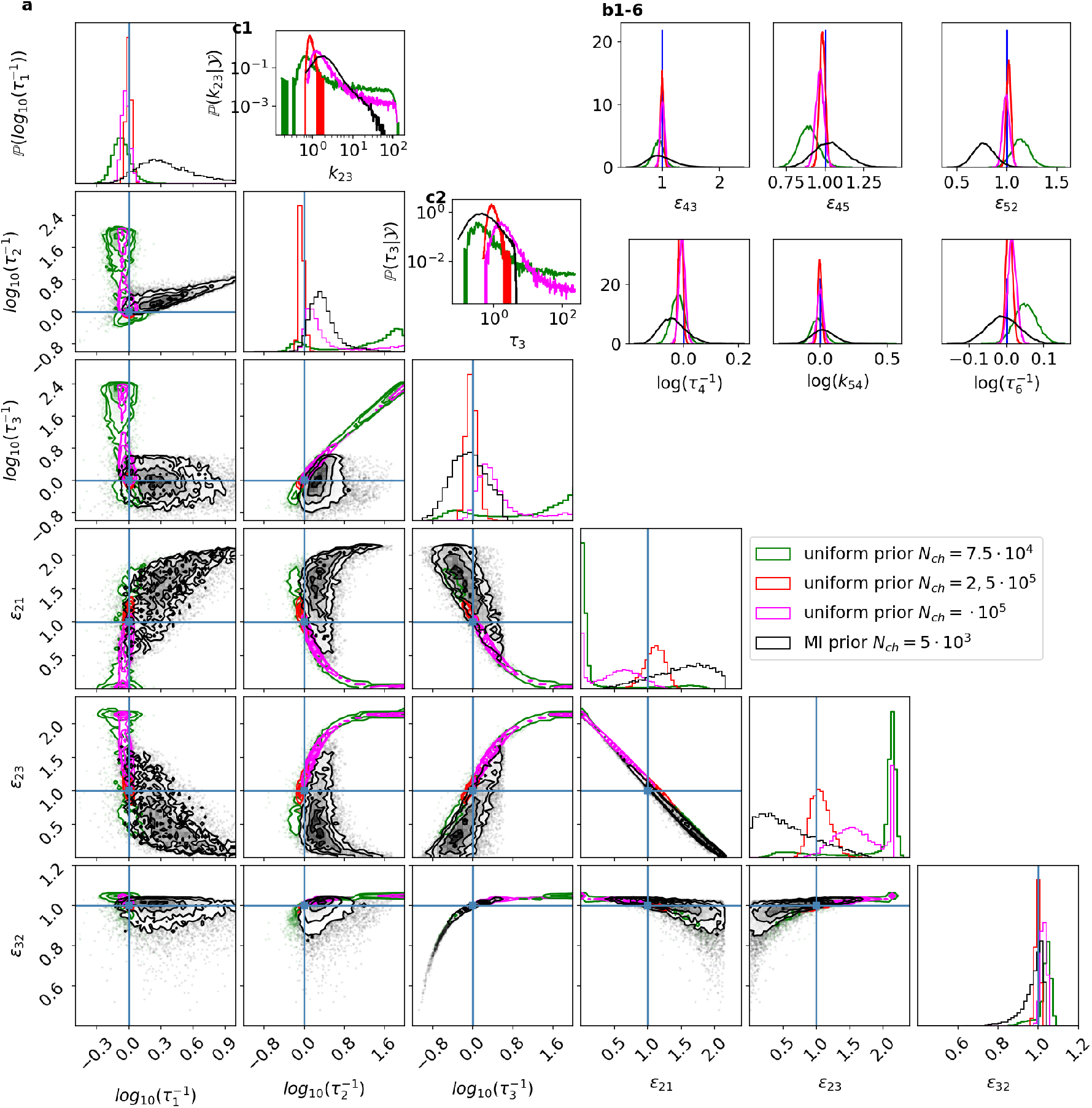
The minimally informative prior increases the data range for a proper posterior for CRN2 by a factor of more than 20. **a**, Four posteriors are compared to show the overall advantage in practical identifiability (*sufficiently proper* posteriors) gained by the minimally informative prior. Note that the colored posteriors are all using higher *N*_ch_ and smaller simulation boxes, which makes it, in principle, easier for these posteriors to hide their practical non-identifiability pathology, resulting in an improper posterior. We suppress all marginal posteriors of somewhat boring parameters. *Boring* in the sense that they deliver more or less Gaussian-shaped marginal posteriors under all tested conditions. The black posterior derived from *N*_ch_ = 5 · 10^3^ is equipped with the minimally informative *p*(**K**). The colored posteriors use a uniform *p*(**K**). They used data sets with *N*_ch_ = [2 · 10^4^ (green), 10^5^ (magenta), 2.5 · 10^5^ (red)] equipped with the uniform *p*(**K**). The flat likelihood for *k*_32_ and *τ*_3_ creates an exponential increase for the posterior of *p*(log_10_(*k*_32_) | 𝒴 _*T*_) and *p*(log_10_(*τ*_3_) | 𝒴 _*T*_). Note that the simulation box for the posterior samples of *τ*_2_ ∈ [10^*−*2^, 10^6.5^] updated from the minimal information is more than an order of magnitude larger than the range of the simulation box used when the uniform *p*(**K**) is employed. The posteriors derived from the uniform *p*(**K**) are sampled by *k*_*ij*_ and then mapped to the (***τ***, ***ϵ***)-space **b**, prior that is used to allow only for minimally violated microscopic reversibility.

For the uniform prior, 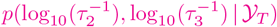, derived from *N*_ch_ = 10^5^ per patch, still displays the practical non-identifiability problem visually. Hence, 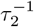 and 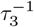 should be considered unidentified, based on the herein-employed heuristic criterium (see, discussion around Eq. 9) to define *p*(**K** | 𝒴 _*T*_) as *sufficiently proper* and the parameter identified if the existing practical non-identifiability pathologies are not detected *visually*. However, 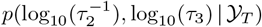 of the data set with 20 times smaller *N*_ch_ and based on the minimally informative *p*(**K**) does not display the practical non-identifiability problem. In contrast, for the next smaller data set (not shown here), with *N*_ch_ = 2 · 10^3^ also 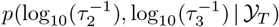 derived from the minimally informative *p*(**K**) starts to display the practical non-identifiability problems visually on a linear scale. For *N*_ch_ = 2.5 ° 10^5^, *p*(log_10_(*k*_32_), log_10_(*τ*_3_) | 𝒴 _*T*_) updated from the uniform *p*(**K**) does not display the practical non-identifiability problem visually anymore. Hence, the minimally informative *p*(**K**) increases the range of acceptable data for this CRN at roughly a 50 (at least a 20) fold, likely more for this data-generating process. Note that 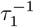 is little influenced by the practical non-identifiability problem in 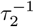 and 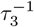. Just as naively expected, the uniform prior based 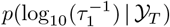 are more narrow and closer to 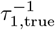 as these posteriors originate from higher quality data. Note that 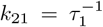 holds because only one Markov transition leaves *C*_1_ (CRN2, Fig. 2 **d**)

### 1. Finding a needle in an *almost* infinite space: The minimally informative prior helps the sampler to converge

To have a decently converging HMC sampler, we used for the sampling of *p*(**K** | 𝒴 _*T*_) based on uniform *p*(**K**) an about ten times smaller sampling box in the direction of *k*_32_ and *τ*_3_. It is plausible that ridges with a bimodal structure, *p*(log_10_(*k*_32_), log_10_(*τ*_3_) | 𝒴 _*T*_) (green and magenta Fig. 15), cause the convergence problems. The larger the simulation box in the directions of *k*_32_ and *τ*_3_ the more would the peak around the true value and the ridge between the peaks lose its probability mass until an independently started chain [104–106] typically explores only one concentration of probability mass or the other but not both. At some size of the sampling box, the peak of the posterior around the true **K** will be like a needle in the vast constant parts of *p*(**K** | 𝒴 _*T*_) and not be found by the sampler, unless one changes from uniform to a minimally informative prior, giving the constant parts of the posterior the slope of the minimally informative prior. Inevitably, the second peak exists due to the log_10_ transformation of the flat posterior in the linear space to the posterior on the log space. The marginal posterior on this part of ***θ*** without the logarithmic transformation is flat (Fig. 15 **c**.**1, c**.**2**). Note that a comparison with a larger sampling box would disfavor the uniform prior even more because of the practical non-identifiability problem. The relative positive effect on the range of acceptable data quality of the minimally informative prior grows with the use of larger sampling boxes.

As mentioned above, the limits for a proper enough *p*(**K** | 𝒴 _*T*_) based on the uniform prior is somewhere between *N*_ch_ = 10^5^ and *N*_ch_ = 2.5 · 10^5^. Unfortunately, using the visual criterium based on the uniform prior requires placing the sampling box limits with a lot of luck around the peak of *p*(**K** | 𝒴 _*T*_), which might be possible if before a ***θ***_ML_ is performed. However, this is not helpful when inferencing HMMs from actual experiments with the typical data amount typically much below *N*_ch_ = 10^4^ and data imperfections.

We made this compromise of sampling the posterior based on the uniform prior in a smaller sampling box to be able to get a reasonably converging HMC sampler in the sense that four independently started sampling chains result in the same *p*(**K** | 𝒴 _*T*_), as judged by the Gelman-Rubin [104, 107] statistic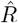. For *N*_ch_ = 2 · 10^4^ and the uniform *p*(**K**) the sample traces (green posterior) have the desired 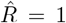 calculated over the four independently started sampling chains for all parameters. However for *N*_ch_ = 10^5^ and *N*_ch_ = 2.5 · 10^5^ one chain out of 4 chains starts to show a different sample trace. In principle, calculating the posterior from only one independent chain would be enough if one would know that this chain represents the typical set of the posterior [73] and has a large enough effective sample size [106]. If we only calculate the posteriors from the three seemingly convergent chains, the posteriors (magenta and red posterior) are well within the predicted probability mass of the posterior (*N*_ch_ = 2 · 10^4^, green). That is the expected behavior of the posterior of a much larger amount of data. Calculating 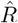 from the three similar chains 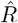, we obtain the demanded 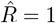. The minimally informative prior alleviates these challenges. Note, that because for the simulated data we know the ground truth of the parameters such that we have a third independent criterion judging the remaining sampling chains. Does the posterior derived from them cover the true values.

